# Cryo-EM Structures of Amyloid-β Fibrils from Alzheimer’s Disease Mouse Models

**DOI:** 10.1101/2023.03.30.534981

**Authors:** Mara Zielinski, Fernanda S. Peralta Reyes, Lothar Gremer, Sarah Schemmert, Benedikt Frieg, Antje Willuweit, Lili Donner, Margitta Elvers, Lars N. G. Nilsson, Stina Syvänen, Dag Sehlin, Martin Ingelsson, Dieter Willbold, Gunnar F. Schröder

## Abstract

The development of novel drugs for Alzheimer’s disease has proven difficult, with a high failure rate in clinical trials. Typically, transgenic mice displaying amyloid-β peptide brain pathology are used to develop therapeutic options and to test their efficacy in preclinical studies. However, the properties of Aβ in such mice have not been systematically compared to Aβ from the patient brains. Here, we determined the structures of nine *ex vivo* Aβ fibrils from six different mouse models by cryo-EM. We found novel Aβ fibril structures in the APP/PS1, ARTE10, and tg-SwDI models, whereas the human familial type II fibril fold was found in the ARTE10, tg-APP_Swe_, and APP23 models. The tg-APP_ArcSwe_ mice showed an Aβ fibril whose structure resembles the human sporadic type I fibril. These structural elucidations are key to the selection of adequate mouse models for the development of novel plaque-targeting therapeutics and PET imaging tracers.

**One Sentence Summary:** Cryo-EM structures of Aβ fibrils extracted from brains of mouse models used for Alzheimer’s disease preclinical research are presented.

## Main Text

Alzheimer’s disease (AD) is the most common form of dementia and is neuropathologically defined by the presence of extracellular plaques containing amyloid-β (Aβ) in the brain parenchyma and intraneuronal neurofibrillary tangles containing phosphorylated tau [1]–[4]. In the amyloidogenic pathway, Aβ is sequentially cleaved from the amyloid precursor protein (APP) by β- and ɣ-secretases [5], [6]. Typically, depending on ɣ-secretase cleavage, monomers between 37 and 43 residues in length are generated, however, the most abundant peptides are 40 (Aβ40) and 42 (Aβ42) residues in length [7]. These monomers tend to aggregate into insoluble fibrils, the structure of which has been extensively studied *in vitro* by cryogenic-electron microscopy (cryo-EM) and solid-state nuclear magnetic resonance (NMR) spectroscopy, revealing a spectrum of different polymorphs [8]–[14]. However, these fibrils are structurally different from both Aβ40 and Aβ42 fibrils derived from AD brain tissue by seeded growth [15], [16] as well as Aβ40 and Aβ42 fibrils extracted from AD meninges [17] and parenchyma [18], respectively. Yang and colleagues determined two human fibril polymorphs: ‘type I filaments’, which are mostly associated with sporadic AD (sAD) and ‘type II filaments’ observed in familial AD (fAD) and other neurodegenerative disorders [18]. Animal models are an important tool to study the pathogenesis of AD and to conduct preclinical testing of novel therapeutics [19]. Commonly used animal models are transgenic mice that mimic different clinical characteristics of the disease [20]. The structures of Aβ fibrils extracted from two different mouse models, the knock-in APP^NL-G-F^ and the knock-in APP^NL-F^, were recently determined by cryo-EM [18], [21], [22]. While the APP^NL-F^ Aβ42 fibril resembles the human type II Aβ42 fold, the fold of the APP^NL-G-F^ Aβ42(E22G) fibrils differs from fibrils extracted from human brain.

To date, there is no cure for AD, but aggregated Aβ is a common target for drug development [23]. One of the most successful treatments so far is the Aβ-directed antibody lecanemab which was developed primarily against intermediately sized soluble Aβ aggregates, in particular, Aβ oligomers and protofibrils [24], [25]. The structure of these Aβ aggregates remains elusive, but a recent study [26] showed interactions between lecanemab and Aβ fibrils that were present in “ultracentrifugal supernatants of aqueous extracts from AD brains”. Nevertheless, developing novel drugs for AD has been challenging, with a drug development failure rate of almost 100% [27]. Structural differences between human and murine Aβ fibrils may explain why positron emission tomography (PET) imaging tracers fail to detect Aβ deposits in some patients (e.g., patients with the Arctic mutation) [28]. Additionally, it might help to understand why fibril-targeting drug candidates show efficacy when tested in mouse models but then fail to show the desired effect in clinical trials.

Here, we extracted Aβ fibrils from the brains of six different mouse models based on a previously described sarkosyl extraction method [18] and determined their structure by cryo-EM (**Fig. 1, Fig. S1, Fig. S2, Fig. S3, Table S1**). The investigated mouse models, APP/PS1, ARTE10, tg-SwDI, tg-APP_Swe_, APP23, and tg-APP_ArcSwe_ are all commonly used lines in AD research and drug development.

**Figure 1:**
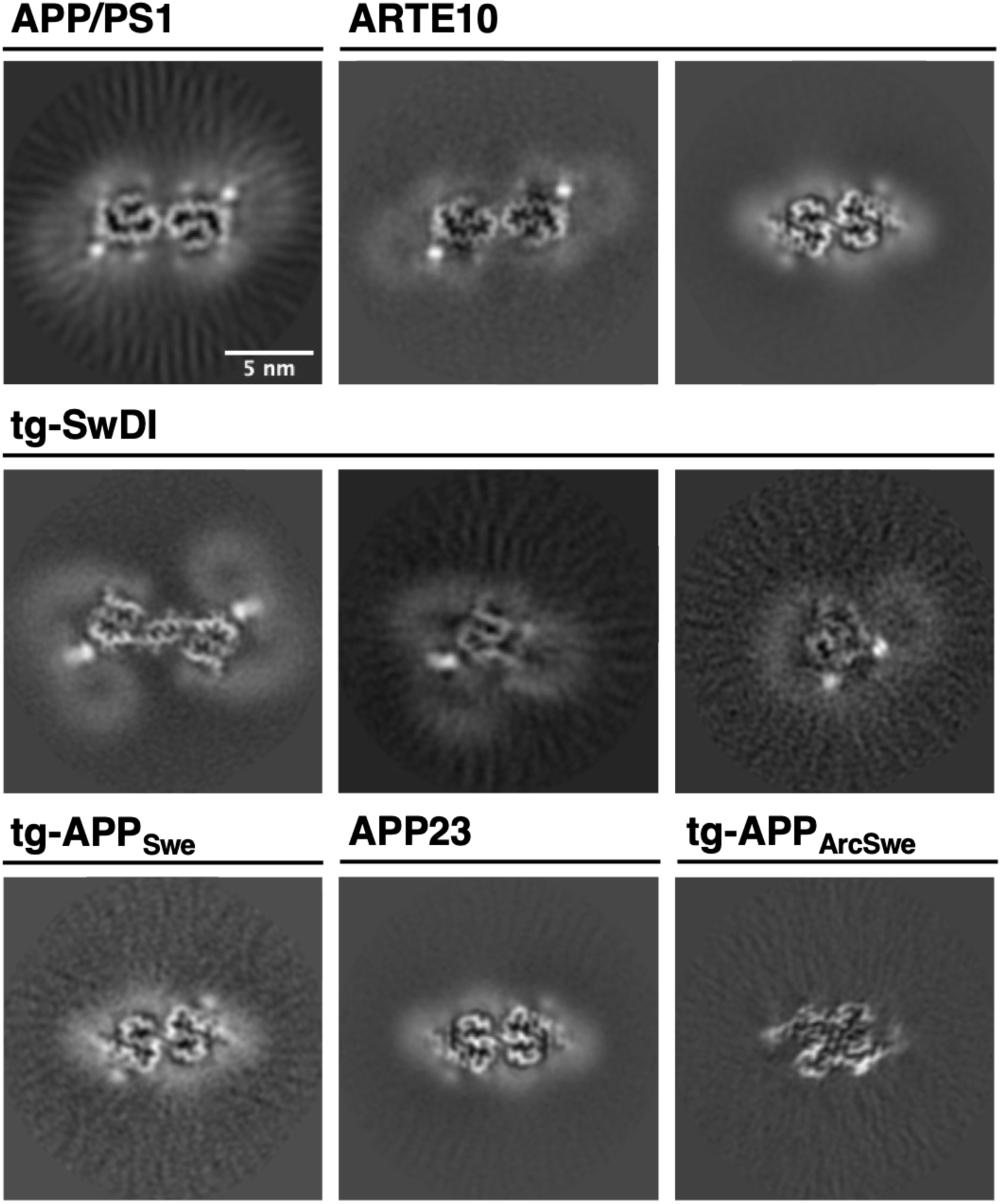
Cryo-EM reconstructions of Aβ fibrils extracted from APP/PS1, ARTE10, tg-SwDI, tg-APP_Swe_, APP23, and tg-APP_ArcSwe_ mouse brain tissue. For every reconstructed fibril, a cross-section through the reconstructed density map is shown. The scale bar in the top left panel applies to all shown panels. From upper left to lower right: murine type III (APP/PS1), murine type III (ARTE10), murine type II (ARTE10), DI1, DI2, DI3, murine type II (tg-APP_Swe_), murine type II (APP23) and murine_Arc_ type I.

### Murine type III Aβ fibrils from APP/PS1 and ARTE10 mouse brains

For the APP/PS1 mice, we observed only one fibril type made of two identical LS-shaped protofilaments related by a C2 symmetry (**Fig. 1 and Fig. 2, A and B**). This fibril type, which we call the murine type III Aβ fold in the following, was also found in ARTE10 mouse brain (**Fig 2C**) and accounts for 4% of all reconstructed ARTE10 fibrils (**Table S2**). Murine type III fibrils were determined to a resolution of 3.5 Å and 3.3 Å for APP/PS1 mice and ARTE10 mice, respectively (**Table S3 and Fig. S4, A and B**). For APP/PS1 murine type III Aβ42 fibrils, atomic model building was possible for the ordered core from residues G9-A42 (**Fig. 2, A and B**). Interestingly, in contrast to ARTE10 murine type II, the reconstructed density of ARTE10 murine type III suggests that the fibril is composed of Aβ40 rather than Aβ42. Accordingly, an atomic model was built from residue G9-V40 (**Fig. 2, A and C**). The N-terminal L-turn involves residues Y10-F19 and is mainly stabilized by one hydrophobic cluster composed of Y10, V12, L17, F10, L34 and V36 (**Fig. 2, B and C and Fig. S5, A and B**). The S-turn, which involves residues F20-V40/A42, is stabilized by two hydrophobic clusters: in the first half of the S-turn between F20-K28 involving A21, V24 and I31 and the C-terminal second half of the S-turn involving A30, I32, M35, V40 (and A42). The protofilament interface involving residues D23-K28 of murine type III fibrils is stabilized by symmetric salt bridges between D23 and K28 of the opposing subunits.

**Figure 2:**
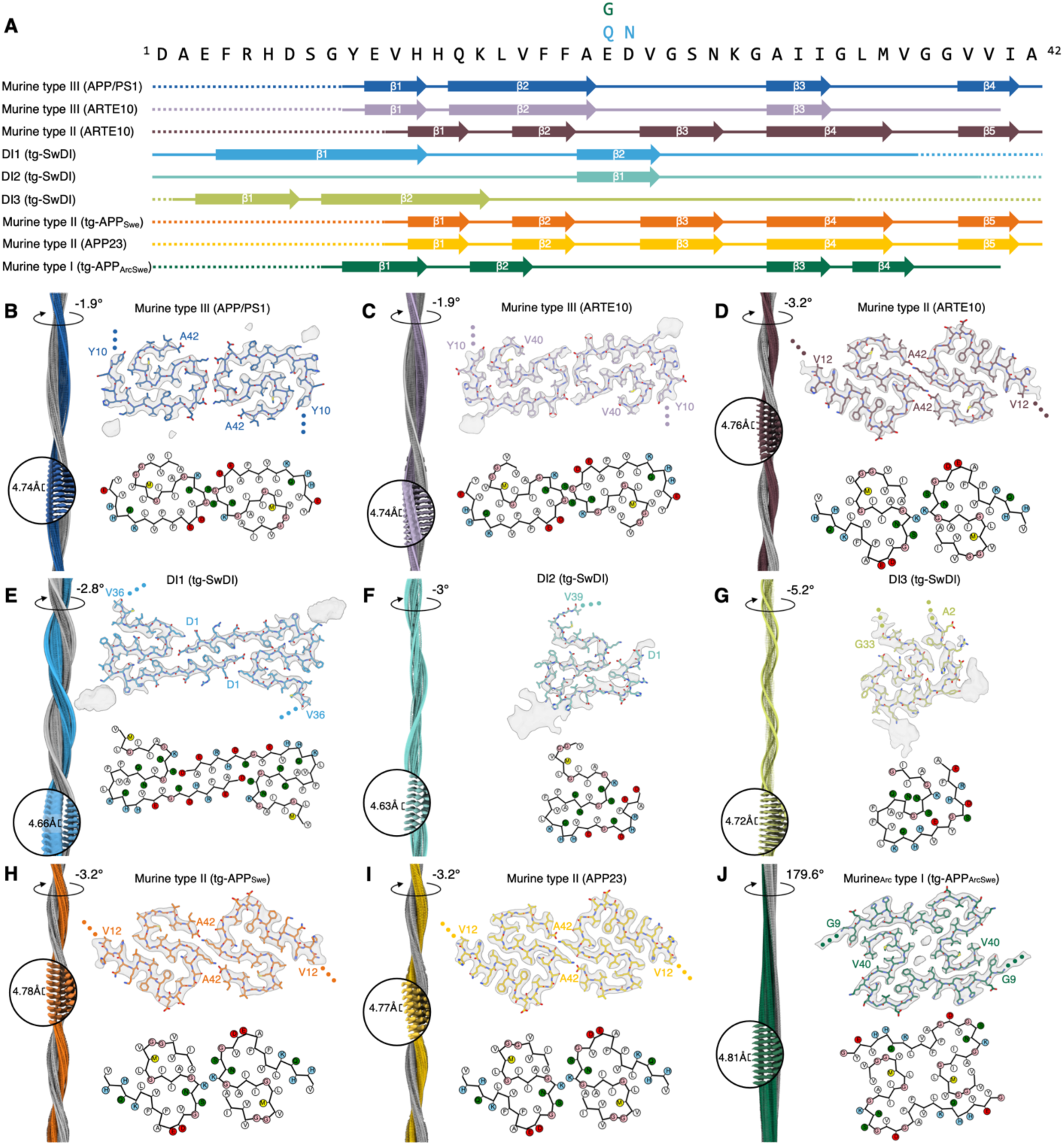
Overview of all murine Aβ fibril structures. (A) Amino acid sequence of Aβ42. The sequence contains the following mutations for tg-SwDI: E22Q and D23N; and for tg-APP_ArcSwe_: E22G. Solid lines indicate for which part of the sequence atomic model building was possible (accordingly, dotted lines represent parts of the sequence that were not modelled). Arrows indicate β-strands. Panels (B-J) consist of: the reconstructed cryo-EM density along the helical axis and a close-up with labels denoting the helical twist and rise (left); the cryo-EM density map (in transparent gray) with the corresponding atomic model (top right); a schematic of the fold, produced with atom2svg.py [51] (red: acidic side chain; blue: basic side chain; green: hydrophilic side chain; white: hydrophobic side chain; pink: glycine; yellow: sulfur containing) (bottom right). Cryo-EM structure of (B) murine type III Aβ42 fibrils from APP/PS1 mouse brain, (C) murine type III Aβ40 fibrils from ARTE10 mouse brain, (D) murine type II Aβ42 fibrils from ARTE10 mouse brain, (E) DI1 Aβ fibrils from tg-SwDI mouse brain, (F) DI2 Aβ fibrils from tg-SwDI mouse brain, (G) DI3 Aβ fibrils from tg-SwDI mouse brain, (H) murine type II Aβ42 fibrils from tg-APP_Swe_ mouse brain, (I) murine type II Aβ42 fibrils from APP23 mouse brain, and (J) murine_Arc_ type I Aβ40 fibrils from tg-APP_ArcSwe_ mouse brain.

Interestingly, similarities can be observed between murine type III fibrils and human Arctic (E693G, E22G in Aβ) Aβ filaments [21] (**Fig. 3A**). The human Arctic Aβ filament shows two distinct protofilaments A and B, with each being present twice in the four-protofilament fibril. The main chain trace of murine type III fibrils resembles one protofilament A-B pair of human Arctic Aβ-filaments. The structures including side chain orientations are identical between E22/G22 and the C-terminus of Aβ, leading to the same solvent exposed residues. Moreover, in both cases the interface is stabilized by salt bridges between D23 and K28. The largest deviation between the two structures can be found in the orientation of the side chains in the N-terminal part up to the single point mutation site (E22G).

**Figure 3:**
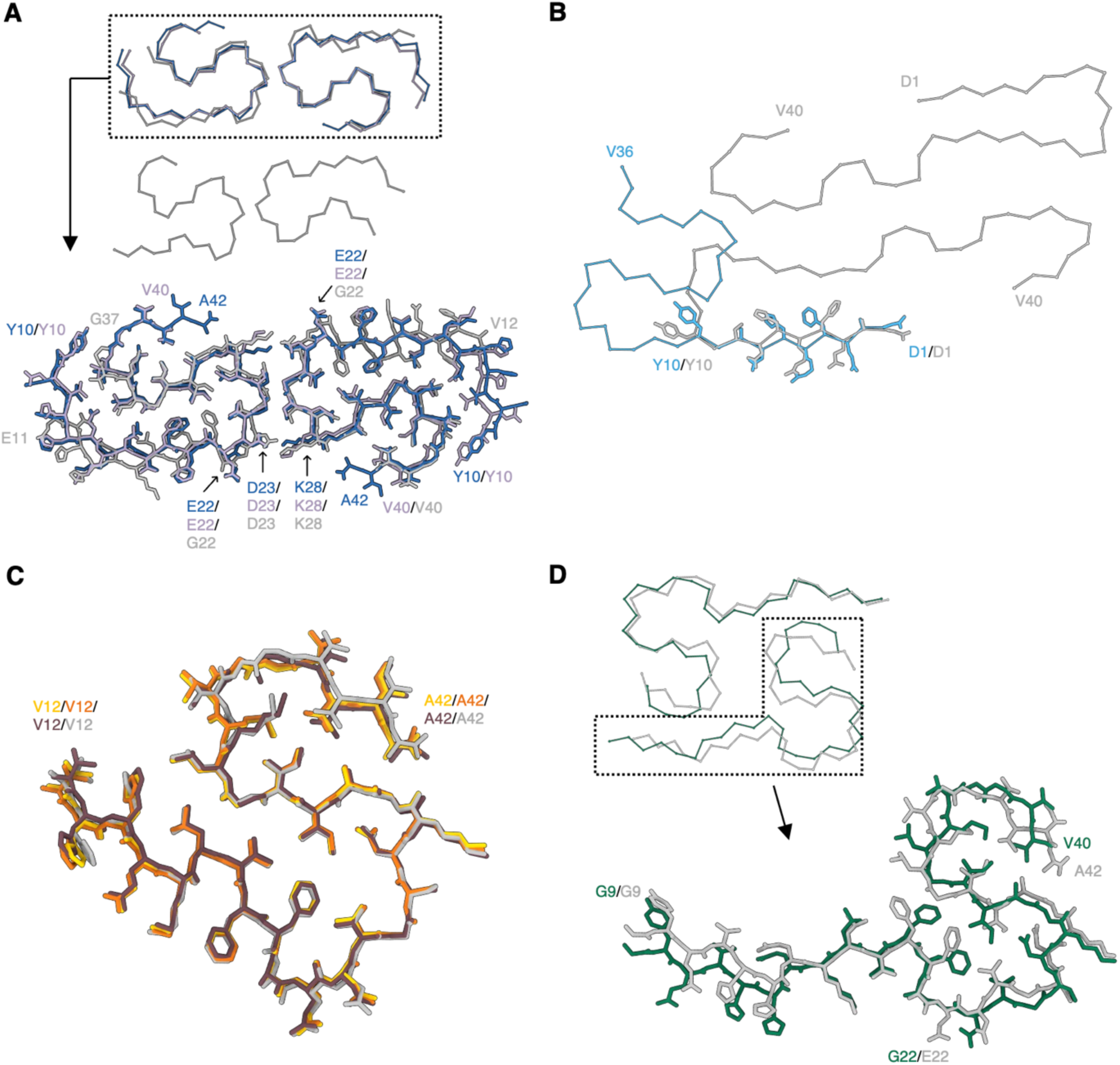
Comparison of brain-derived murine Aβ fibrils to brain-derived human extracted Aβ fibrils. (A) Comparison of murine type III Aβ fibrils (blue: APP/PS1; lavender: ARTE10) with the cryo-EM structure of human brain extracted Aß filaments with the E693G (E22G) mutation (gray, PDB code: 8BG0), (B) Comparison of the DI1 Aβ fibril (light blue) with the cryo-EM structure of Aβ40 fibrils extracted from the meninges of human AD brain tissue (gray, PDB code: 6SHS), (C) Comparison of the APP23 (yellow), tg-APP_Swe_ (orange) and ARTE10 (burgundy) Aβ42 fibril fold with human type II Aβ42 filaments (gray, PDB code: 7Q4M), (D) Comparison of the murine_Arc_ type I (green) Aβ40 fibril fold with human type I Aβ42 filament fold (gray, PDB code: 7Q4B).

### Novel Aβ folds from tg-SwDI mouse brain

For tg-SwDI mice, which harbor the Dutch (E22Q) and Iowa (D23N) mutations within the Aβ sequence, we observed three different polymorphs (**Fig. 1 and Fig. 2, A, E, F, and G**). The most dominant polymorph, which we call DI1, accounts for 41% of all fibrils and reveals a symmetric dimer (**Fig. 2, A and E and Table S2**). The other two polymorphs, labeled DI2 and DI3, consist of a single protofilament and account for 32% and 27%, respectively (**Fig. 2, A, F, and G and Table S2**).

The 3.3 Å map of DI1 was used to build an atomic model of the ordered core between D1 and V36 (**Fig. 2, A and E and Table S3 and Fig. S4D**). The two S-shaped protofilaments of DI1 are connected via the extended N-terminus (**Fig. 2E**). A hydrogen bond between D1 and S26 and a salt bridge between E3 and K28 of the opposing protofilament stabilize the interface between the two subunits. A hydrogen bond between Q15 and N23 stabilizes the first half of the S-turn. This turn is further stabilized by a hydrogen bond between Y10 and a backbone oxygen at N23^i-1^ in the adjacent layer within the same protofilament (denoted by the index i-1).

For DI2 fibrils an atomic model of residues D1-V39 could be built into the 4.2 Å map (**Fig. 2, A and F and Table S3 and Fig. S4E**). Except for the N-terminus, the fold is similar to the fold of DI1 fibrils (**Fig. 2, E and F and Fig. S6A**). The N-terminus is fixed in its position by a salt bridge between D1 and K28^i+2^ as well as a hydrogen bond between E3 and S26^i+2^. The second β-sheet found in DI1 is also present in DI2 (**Fig. 2A**).

For DI3, the atomic model consists of A2-G33 (**Fig.2, A and G**). The overall DI3 fold differs from the DI1 fold and aligns only in residues V24-G33 with the DI2 fold (**Fig. 2, E, F, and G and Fig. S6A**). Therefore, secondary structure assignments differ from DI1 and DI2, showing two β-sheets in the N-terminal domain (**Fig. 2A**). The fold of DI3 fibrils is mainly stabilized by a salt bridge between E11 and K28^i+1^ and a hydrogen bond between E11 and S26^i+1^. The C-terminal kink in the structure around K28 is stabilized by a hydrogen bond between N27 and the carbonyl group at G29. Additionally, a hydrophobic cluster around V18, A21 and V24 stabilizes the overall fold (**Fig. S5F**).

Aβ fibrils extracted from tg-SwDI mouse brain are structurally different from Aβ fibrils extracted from human and APP^NL-F^ mouse brain tissue [18]. However, DI1 fibrils show some structural similarities to Aβ40 fibrils extracted from the meninges of human AD brain (**Fig. 3B**) [17] and from APP^NL-G-F^ mouse [21], [22] (**Fig. S6B**). Remarkably, unlike in most other known Aβ structures, the N-terminus in human Aβ40, APP^NL-G-F^ as well as in DI1, DI2 and DI3 fibrils is ordered. Here, the extended N-terminal arm of DI1 overlays with the N-terminus of the human Aβ40 fibril between residues D1 and Y10 and with the murine Arctic Aβ filament between E3 and S8. Moreover, the orientation of these side chains is identical in all structures, suggesting the same degree of solvent-accessibility of the N-terminus.

Finally, in contrast to wild-type (WT) Aβ42 fibrils, in which negatively charged and solvent exposed residues E22 and D23 induce a kink in the main chain, in DI1 and DI2 fibrils mutant residues Q22 and N23 are interior, in extended conformation (**Fig. 2, E and F**).

### Murine type II Aβ Fibrils from ARTE10, tg-APP_Swe_, and APP23 mouse brains

Aβ fibrils extracted from ARTE10, tg-APP_Swe_, and APP23 mouse brains are composed of Aβ42 and identical to previously described type II filaments extracted from human brain tissue of familial AD [18], therefore here they are referred to as murine type II (**Fig. 1 and Fig.2, A, D, H, and I and Fig. 3C**).

The residues V12-A42 form the ordered core of all three murine type II fibrils, whereas N-terminal residues D1-E11 are likely flexible and therefore not resolved in the high-resolution cryo-EM structure (**Fig.2, A, D, H, and I**). Murine type II fibrils are made of two S-shaped protofilaments, which are related by a C2 symmetry in all three models. Each monomeric subunit is stabilized by two hydrophobic clusters around residues (i) L17, V18, F20, V24, N27, I31, and L34 and (ii) A30, I32, M35, V40, and A42 (**Fig.2, A, D, H, and I and Fig. S5, C, G, and H**). The interface between the two protofilaments is rather small, involving only two symmetric hydrogen bonds between K28 and A42 of the opposing subunit.

Additionally, as previously discussed by Yang et al. [18], murine models that resemble the type II filament fold overlap in their S-shaped domain partially with seeded Aβ40 fibrils extracted from cortical tissue of an AD patient [15] (**Fig. S6C**).

### Aβ Fibrils from tg-APP_ArcSwe_ mouse brain

The sample of Aβ fibrils with the Arctic mutation (E693G, E22G in Aβ) extracted from tg-APP_ArcSwe_ mouse brain tissue consisted of at least three different polymorphs. Structure determination was possible for the most abundant polymorph (**Table S2 and Table S3**), which we call the murine_Arc_ type I Aβ fibril in the following, with a crossover distance of ∼950 Å and a diameter of 90 nm (**Fig. 1**). Murine_Arc_ type I Aβ fibrils consist of two identical S-shaped protofilaments that are related by a pseudo-2_1_ symmetry (**Fig. 2J**). The 3 Å-resolution map of murine_Arc_ type I Aβ fibrils allowed for atomic model building of the ordered core from G9-V40, in agreement with an observed predominance of Arctic Aβ40 fibrils in the sample (**Fig. 2, A and J and Table S3 and Fig. S4I**) [25], [29]. The S-shaped domain is formed by the residues F19-V40, with an associated extended N-terminal arm of G9-V18 that interacts with the C-terminus of the opposing protofilament. The S-shape is stabilized by two hydrophobic clusters around (i) F19, F18, V24, and I31 and (ii) A30, I32, M35, and V40 (**Fig. 2J and Fig. S5I**). The interface between the two protofilaments consists mainly of hydrophobic interactions involving the side chains of Y10, V12, Q15, L17, V36, and V39 at the contact point of the N-terminus of one protofilament and the C-terminus of the opposing protofilament. The fibril center harbors a hydrophobic cavity between the two C-terminal domains of both protofilaments, where two isolated symmetric densities can be observed indicating the presence of additional hydrophobic molecules of unknown identity in the interface.

Murine_Arc_ type I Aβ40 fibrils resemble human type I Aβ42 filaments found in sAD patients for the most part (**Fig. 3D**) [18]. In detail, the solvent-accessible surface is almost identical to human type I Aβ42 filaments, but the C-terminus is slightly shifted, likely caused by the bound molecules in the hydrophobic fibril cavity. Aβ filaments from APP^NL-G-F^ knock-in mice also carry the Arctic mutation [21], [22], and although APP^NL-G-F^ Aβ filaments and murine_Arc_ type I Aβ40 fibrils share a common substructure between residues L17-V36, the overall fold and the resulting arrangement of the two protofilaments differs (**Fig. S6D**). Additionally, and in contrast to APP^NL-G-F^ Aβ filaments, in which the mutant residue G22 is hidden in the protofilament interface, the G22 in murine_Arc_ type I Aβ fibrils is exposed to solvents similarly to E22 in human type I filaments. To date, there is no other murine model known that contains predominantly Aβ fibrils that mimic the human type I fold.

The only *in vitro* preparation that resembles the Aβ40 murine_Arc_ type I fold to a large extent is the NMR structure of Aβ40 fibrils with the Osaka mutation (E693Δ, E22Δ in Aβ) [30] (**Fig. S6E**). In both cases, the mutation (or deletion) of the acidic residue E22 in Aβ40 results in fibrils highly similar to sAD fibrils. Additionally, it was previously shown that Aβ42 fibrils that also harbor the Arctic mutation extracted from knock-in APP^NL-G-F^ mice brain do not resemble human type I filaments [21], [22]. As discussed for murine type II fibrils, the murine_Arc_ type I fibrils also partially overlap with the cryo-EM structure of brain homogenate seeded Aβ40 [15] (**Fig. S6C**).

### Additional densities in murine Aβ fibrils

We have observed additional densities on the surface of all murine fibrils. Strong, localized densities can be observed close to K16 in murine type III, DI1, DI2, DI3, and murine type II Aβ fibrils (**Fig.1 and Fig. 2 and Fig. S7, A, B, and C**). Moreover, murine type III fibrils show a smaller density close to F20 and E22 (**Fig. 7D**) and in DI3 fibrils, an additional density can be found next to Y10. Similar densities bound to K16 were previously described for APP^NL-G-F^ mice but not for AD patients [18], [21]. The observed additional density might be related to bound co-factors or post-translational modifications such as ubiquitination, as it was previously described for tau filaments [31]. Weak, micelle-like density of unknown origin bound to the fibril surface is visible in tg-SwDI Aβ fibrils (**Fig.1 and Fig. S7B**), reminiscent of previously described densities on the surface of alpha-synuclein fibrils [32].

## Discussion

### Here we extracted Aβ fibrils from six different transgenic mouse models and determined their structure using cryo-EM

We observed novel Aβ fibril folds in brain extracts from three mouse models (APP/PS1, ARTE10, and tg-SwDI). Although murine type III Aβ fibrils extracted from APP/PS1 and ARTE10 mice showed some similarities to the human Arctic Aβ filament [21], their fold has not yet been observed in AD patients. The tg-SwDI model harbors a triple mutation [33], [34] that cannot be found in patients, however, these have been observed individually in early-onset AD families. DI1, DI2, and DI3 fibril structures differ from human type I and type II filaments [18] and, consequently, tg-SwDI might not be an appropriate model for neither sAD nor fAD. However, tg-SwDI is also used to study cerebral amyloid angiopathy (CAA) and indeed, the N-terminus of DI1 fibrils is identical to that of a previously described human Aβ40 polymorph obtained from vascular deposits in the brain meninges associated with CAA [17]. Therefore, tg-SwDI mice could indeed be appropriate to study CAA.

Additionally, Aβ fibrils from three mouse models (ARTE10, tg-APP_Swe_, and APP23) resemble the human fAD type II filament fold and are therefore, together with the previously described knock-in APP^NL-F^ model [18], possible candidates for therapeutic research focused on fAD. For example, the APP23 line underwent treatment with different aducanumab analogs, showing positive results such as reduction of total plaque area and improvement in spatial memory [35] Therefore, preclinical testing with aducanumab might have been predictive for efficacy in fAD, but not well tested for sAD.

Aβ fibrils extracted from tg-APP_ArcSwe_ mouse are almost identical to human type I filaments [18] and are therefore here referred to as murine_Arc_ type I fibrils. Human type I filaments are mainly found in sporadic AD, which accounts for more than 95% of all such cases [36]. However, in contrast to human type I filaments, which are composed of wildtype Aβ42, murine_Arc_ type I fibrils consist of the E22G variant of Aβ40. Moreover, murine_Arc_ type I fibrils show two additional densities in the protofilament interface. The Arctic mutation is also present in knock-in APP^NL-G-F^ mice, but their Aβ(E22G) fibril structure differs from murine_Arc_ type I fibrils and human type I filaments [18], [21], [22]. In therapeutic research, tg-APP_ArcSwe_ mice were treated with the mAb158 monoclonal antibody promoting protofibril clearance [37]–[39]. A humanized version of the mAb158 antibody, named as BAN2401 but most commonly known as lecanemab, showed deceleration of cognitive decline and reduction of amyloid plaque burden in the brain of AD patients [40]–[43]. So far, our murine_Arc_ type I structure is the only fibril structure that resembles the human type I filament fold and therefore, tg-APP_ArcSwe_ might be a valuable model to predict which drug candidate will show efficacy in sAD. The structural similarity between murine_Arc_ type I filaments and human sporadic type I filaments might explain the clinical success of lecanemab [43], especially considering that it binds not only to intermediately sized soluble aggregates, but also to “diffusible Aβ fibrils” whose structure is identical to that of Aβ fibrils found in insoluble plaques [25], [26], [40].

[^11^C] Pittsburgh compound B (PiB) and later developed fluorine-18 (^18^F) radiolabeled analogues are commonly used PET tracers to detect AD pathology in the living brain. A positive PET scan has served as an inclusion criterion in clinical studies of antibodies such as aducanumab and lecanemab and further, a reduction in the PET signal intensity has been interpreted as successful removal of brain amyloid plaque, and thus, included as a secondary endpoint in the clinical trials. Interestingly, it was observed that patients with the human Arctic mutation are “PET-negative” [28] in spite of having massive Aβ-deposition *post mortem* [44], [45]. However, in line with our result showing that the murine_Arc_ type I structure found in the tg-APP_ArcSwe_ model resembles the human type I filament fold found in human sAD, PET imaging performed in the tg-APP_ArcSwe_ mouse model with [^11^C]PiB visualizes amyloid pathology [25]. Likewise, [^11^C]PiB also works effectively in the ARTE10, tg-APP_Swe_, and APP23 mouse models, that show an Aβ fibril fold similar to human type II filaments, as long as the studied mice exhibit high total brain Aβ levels. Yet, [^11^C]PiB does not work effectively in every mouse model, as is the case for the APP/PS1 model and interestingly, that model contains murine type III fibrils which are similar to those of human Arctic brain which is also PiB-negative [28]. Thus, it is believed that the ability of [^11^C] PiB to detect pathology depends on differences in the structure of amyloid plaques and Aβ fibrils therein [46]–[49]. For example, it has been shown that the tg-APP_ArcSwe_ model exhibits higher [^11^C]PiB binding that the APP^NL-G-F^ model [50], whose purified Aβ structures differ from human type I and type II filaments. Furthermore, a recent study that used the ^18^F-labeled amyloid PET tracer florbetaben (FBB) to directly compare the APP/PS1 and the ARTE10 mouse models showed that the ARTE10 mice are more suitable for amyloid-PET due to their congophilic dense-cored plaques and overall higher plaque load compared to the APP/PS1 mouse model [49].

Therapeutic approaches that succeeded in animals and failed to produce positive outcomes in humans may have overlooked the possibility that animal models might not contain the relevant molecular drug targets for sAD as they do not necessarily present the same folds and surfaces. Considering that most of AD patients have a sporadic background, one can infer that this might be one important reason why the failure rate of clinical trials has been so high [27]. Structural studies of Aβ fibrils from animal models and their comparison to human Aβ fibrils, as presented in this work, reveal the molecular targets, and may help us identify the most adequate animal model for the development of novel AD treatments and PET tracers targeting amyloid deposits.

## Acknowledgments

MZ and GFS. gratefully acknowledge the electron microscopy training, imaging and access time granted by the life science EM facility of the Ernst-Ruska Centre at Forschungszentrum Jülich. MZ, BF and GFS. are grateful for the computing time provided by Forschungszentrum Jülich on the supercomputer JURECA/JURECA-DC at Jülich Supercomputing Center (JSC). The ARTE10 mouse line was a generous gift from Taconic Biosciences Inc. GFS acknowledges support from Alzheimer Forschung Initiative e.V. SaS received funding from the German Federal Ministry for Education and Research (project number 16LW028). MI has received grants from the Swedish Research Council (2021-02793). SS and DS acknowledge funding from the Swedish Research council (2021-01083 and 2021-03524), Alzheimerfonden and Hjärnfonden. D.W. was supported by ‘‘Portfolio Drug Research’’ of the ‘‘Impuls und Vernetzungs-Fonds der Helmholtzgemeinschaft.’’

## Author Contributions

Conceptualization: LG, GFS

Organisation of breeding and characterization of mouse tissue: SaS, SS, DS, MI, LD, ME, AW, LNGN

Extraction of Aβ fibrils: FSPR

Immunogold labelling: MZ, FSPR

Cryo-EM grid preparation and data collection: MZ

Image processing, reconstruction, and model building: MZ, BF, GFS

Visualization: MZ, FSPR

Supervision: LG, DW, GFS

Writing - original draft: MZ, FSPR, LG, GFS

Writing - review & editing: All Authors

## Data and materials availability

Cryo-EM maps have been deposited to the Electron Microscopy Data Bank (EMDB) and to the Protein Data Bank (PDB) under the following accession numbers: EMD-16944 (PDB ID: 8OL3) for murine type III Aβ42 fibrils from APP/PS1, EMD-16960 (PDB ID: 8OLO) for murine type III Aβ40 fibrils from ARTE10, EMD-16949 (PDB ID: 8OL5) for murine type II Aβ42 fibrils from ARTE10, EMD-16959 (PDB ID: 8OLN) for DI1 Aβ fibrils from tg-SwDI, EMD-16957 (PDB ID: 8OLG) for DI2 Aβ fibrils from tg-SwDI, EMD-16961 (PDB ID: 8OLQ) for DI3 Aβ fibrils from tg-SwDI, EMD-16952 (PDB ID: 8OL6) for murine type II Aβ42 fibrils from tg-APP_Swe_, EMD-16942 (PDB ID: 8OL2) for murine type II Aβ42 fibrils from APP23, and EMD-16953 (PDB ID: 8OL7) for murine_Arc_ type I Aβ40 fibrils from tg-APP_ArcSwe_.

## Supplementary materials and methods

### Materials and Methods

#### Animals

In the present study, the following mouse lines were used for experimentation including immunohistochemistry, negative stain sample screening, immunogold negative stain, and cryo-EM:

APP/PS1 (APPswe/PSEN1dE) (heterozygous; n=4 (male = 3; female = 1); age: 27–33 months old) on a C57BL/6;C3H background (strain name B6.Cg-Tg(APPswe,PSEN1dE9)85Dbo/Mmjax) are well described in terms of their behavioral and pathological characteristics [52]–[54]. Depending on the used protocol, APP/PS1 mice develop (contextual and spatial) cognitive deficits by seven months of age. Aβ plaques can be detected by six months of age in hippocampus and cortex, followed by a pronounced gliosis. Abundant Aβ plaques and gliosis are prominent with twelve months of age. Four heterozygous APP/PS1 mice brains were used in this study.

ARTE10 (homozygous; n= 1 (female); age= 24 months old) mouse on a C57Bl/6 background (strain name B6.CBA-Tg(Thy1-PSEN1*M146V,-APP*Swe)10Arte) was a generous gift from Taconic Biosciences Inc. (Germantown, NY, USA). The mice express APPswe (APP KM670/671NL) and PS1-M146V under Thy1.1 regulatory sequences, which leads to the development of a progressive plaque pathology and CAA starting around the age of 3 months [55].

Tg-SwDI mice (heterozygous; n = 4 (all male); age: 26–29 months old) on a C57BL/6 background (strain name C57BL/6-Tg(Thy1-APPSwDutIowa)BWevn/Mmjax) were first introduced by Davis et al. in 2004 as a model to study cerebral amyloid angiopathy (CAA) in AD [56], [57]. Cognitive deficits and Aβ plaques with associated gliosis can be detected by three months of age, increasing, and manifesting with age.

APP23 mice (heterozygous; n= 2 (all male); age= 21 months old) are on a C56BL/6 background (strain name B6.Cg-Tg(Thy1-APP)3Somm/J) and have a 7-fold overexpression of mutant human APP_751_ bearing the pathogenic Swedish mutation. Aβ deposit starting at six months of age and increase in size and number with age [52]. APP23 mice also develop CAA [53].

Tg-APP_ArcSwe_ (heterozygous; n= 1 (male); age= 18 months old) and tg-APP_Swe_ (heterozygous; n= 2 (all male); age= 22 months old) are maintained on a C57BL/6 background [58]. Tg-APP_ArcSwe_ mice harbor the Swedish and the Arctic APP mutations and develop plaque pathology starting at around 6 months of age, while Tg-APP_Swe_ mice that harbor the Swedish mutation have a later onset of plaque pathology starting at 10–12 months of age and then increasing with rapidly with age.

APP/PS1, ARTE10, tg-SwDI, APP23 experiments were performed in accordance with the German Law on the protection of animals (TierSchG §§7–9). Breeding of APP/PS1 mice was approved by a local ethics committee [Landesamt für Natur, Umwelt und Verbraucherschutz Nordrhein-Westfalen (LANUV), North Rhine-Westphalia, Germany, Az: 84-02.04.2014.362] before start of the study. APP/PS1 and tg-SwDI mice were (and can be) purchased by the Jackson Lab (JAX MMRRC Stock# 034829 or JAX MMRRC Stock# 034843).The tg-APP_ArcSwe_ and tg-APP_Swe_ mice were bred under the ethical permit 5.8.18-20401/20 approved by the Uppsala County Animal Ethics board. All mice were kept and bred under controlled conditions with 12/12 h light/dark cycle, 54% humidity, a temperature of 22°C as well as food and water *ad libitum*.

#### Brain tissue characterization

Brain tissue from the tg-APP_Swe_, tg-APP_ArcSwe_, and the ARTE10 mouse models has been extensively characterized in previous studies [29], [49], [55], [58]–[60].

The remaining APP/PS1, tg-SwDI and APP23 mouse lines were immunohistochemically stained as follows: In brief, after cervical dislocation, the brains were snap frozen in isopentane and cut in 20 µm sagittal sections with a microtome. Afterwards, the sections were fixed with 4% paraformaldehyde (PFA) in TRIS-buffered saline (TBS) for 10 min at room temperature (RT). The sections were washed three times with 1% Triton in TBS (TBST) for 5 min and further incubated in 70% formic acid for 5 min at RT for antigen retrieval. The sections were again washed with TBST before incubation with primary antibody overnight at 4°C in a humidified chamber (6E10 (BioLegend, Alexa Fluor 594 anti-β-Amyloid, 803018) and 4G8 (BioLegend, 800708, both diluted 1:500 in TBST with 1% bovine serum albumin (BSA)). The day after, the tissue sections were washed with TBST before incubation with the secondary antibody (only 4G8, goat anti-mouse antibody, Alexa Fluor 488, Invitrogen, diluted 1:300 in TBST and 1% BSA) for 1 h at RT. For cell nuclei staining, the sections were washed again with TBST before incubation with DAPI ((4′,6- Diamidin-2-phenylindol) (Merck, Germany)) for 5 min. Subsequently, the sections were washed three times with TBST before mounting (Fluoromount Aqueous Mounting Medium, Sigma-Aldrich, St. Louis, USA). Images were taken with a LMD6000 microscope (Leica Camera, Germany) with a DFC310 FX camera (Leica Camera, Germany).

#### Extraction of Aβ Fibrils

Aβ fibril extraction was essentially based on a published procedure [18]. In brief, non-fixed mouse brain tissue was snap frozen in -80°C cold isopentane and stored at -80°C previous to experimentation. Between 0.4 and 0.6 g of brain tissue was thawed and manually homogenized in 20-times volume (w/v) of extraction buffer (10 mM Tris-HCl, pH 7.5, 0.8 M NaCl, 10% sucrose, 1 mM EGTA) by applying 300 strokes using a Dounce glass tissue grinder. Subsequently, 10% sarkosyl diluted in _d_H_2_O (Sigma-Aldrich) was added to the homogenate to a final sarkosyl concentration of 2% and was thoroughly mixed 30-times by pipetting up and down. After 1h incubation at 37°C, the homogenate was centrifuged at 10.000 x g for 10 min at 4°C and the resulting supernatant was further ultracentrifuged at 100.000 x g for 60 min at 4°C (Beckman Coulter Optima MAX-XP, TLA55 fixed-angle rotor). After removal of the supernatant, extraction buffer (1 ml/g original tissue mass) was added to the pellet and mixed, followed by 5.000 x g centrifugation for 5 min at 4°C. The supernatant was then diluted 3-fold in dilution buffer (50 mM Tris-HCl, pH 7.5, 0.15 M NaCl, 10% sucrose, 0.2% sarkosyl) and ultracentrifuged at 100.000 x g for 30 min at 4°C. The resulting supernatant was discarded and resuspension buffer (20 mM Tris-HCl, pH 7.4, 50 mM NaCl) was added (100 µl/g original tissue mass) to the sarkosyl insoluble Aβ fibril rich pellet. The pellet was used for further negative staining, immunogold labelling and cryo-EM analysis.

We noticed that the fibril extraction protocol was sensitive to changes in temperature, sarkosyl concentration and frequency of homogenization, therefore, the procedure was optimized accordingly.

#### Negative stain Electron Microscopy

2 µl of the final sarkosyl insoluble fraction, consisting of a homogeneous mixture of the final pellet after fibril extraction and resuspension buffer, were applied onto a glow-discharged 300 mesh carbon-coated copper grid (EM Sciences, ECF300-Cu). The sample was incubated for 2 min and carefully blotted off with filter paper. The sample was then washed once with _d_H2O and blotted off immediately. 2 µl of 1% (w/v) uranyl acetate (UrAc) were applied on the top of the grid, following a 1 min incubation. The UrAc was removed with filter paper and the grid was air-dried. TEM images were acquired using a ThermoFisher Scientific Talos 120C at an acceleration voltage of 120 kV. Images were collected on a 4k x 4k Ceta 16M CEMOS camera using Thermo Scientific Velox Software.

#### Immunogold negative stain Electron Microscopy

Immunogold negative-stain grids for electron microscopy were prepared following [61]. In brief, 3 µl of the final pellet containing the extracted Aβ fibrils were placed on a glow-discharged 300 mesh carbon-coated copper grid (EM Sciences, ECF300-CU) for 2 min. The sample was washed once with _d_H20 and placed in blocking buffer for 15 min, following incubation with Nab228 (Sigma-Aldrich) primary antibody diluted in blocking buffer at a concentration of 2 µg/ml for 1-2 h. Furthermore, the grid was washed with washing buffer and was incubated with 6 nm gold-conjugated anti-mouse secondary antibody (diluted 1:20 in blocking buffer, Abcam) for 1 h. The grid was washed five times with washing buffer and three times with _d_H20 before staining with 1% (w/v) uranyl acetate for 1 min. The sample was air-dried, and EM Images were acquired as described above. Immunogold negative stain EM confirmed that the purified fibrils were indeed Aβ fibrils (**Fig. S2**).

#### Cryo-EM Image Acquisition and Data Preprocessing

For cryo-EM imaging, 2-3 µL of Aβ fibril sample from a single mouse brain was applied to holey carbon grids (Quantifoil 1.2/1.3, 300 mesh), blotted with filter paper for 3-5 s and plunge frozen in liquid ethane using a ThermoFisher Scientific Vitrobot Mark IV, set at 95% humidity and 4°C temperature. Data acquisition was performed on a ThermoFisher Scientific Talos Arctica microscope operating at 200 kV using a Gatan Bioquantum K3 detector in counting mode with a Gatan Bio-quantum energy filter with a slid width of 20 eV, and on a ThermoFisher Scientific Titan Krios G4 operating at 300 kV using a Falcon 4 detector in counting mode. The automated collection was directed by EPU data collection software. Further details are given in **Table S1**.

For helical reconstruction of all datasets, gain-corrected movie frames were aligned and summed into single micrographs on-the-fly using Warp [62]. CTF estimation was performed using CTFFIND4.1 [63].

#### Helical Reconstruction

Helical reconstruction was performed using the helical reconstruction methods in RELION [64], [65]. For all datasets, fibrils were picked automatically using crYOLO [66], [67]. Reference-free 2D classification was performed to separate different polymorphs and to discard low quality particle images.

In the recorded dataset of tg-APP_ArcSwe_ at least three polymorphs could be identified and 2D class averages suggest that all polymorphs have a pronounced unstructured region. Indeed, even after multiple 3D classification and refinement runs, we only obtained class averages revealing unstructured regions at the fibril periphery. Therefore, we performed a masked classification with residual signal subtraction [68] and removed the unstructured regions from the reconstruction.

For ARTE10 murine type II, ARTE10 murine type III, DI2, DI3, tg-APP_Swe_ murine type II, APP23 murine type II and murine_Arc_ type I, a featureless cylinder was used as initial 3D reference. For APP/PS1 murine type III and DI1, an initial 3D reference was computed *de novo* from multiple 2D class averages assuming a helical rise of 4.75 Å and a twist value calculated from the crossover-distance of each fibril observed from larger box 2D class averages [69]. Cylinders were initially low-pass filtered to 40 Å, reconstructed *de novo* initial models were low-pass filtered to 8-10 Å depending on their quality. 3D classification was used to obtain a homogeneous high-quality subset of particles for each fibril type. 3D auto-refinement and subsequent post-processing was performed to fibril the final maps and to calculate the resolution according to gold-standard Fourier Shell Correlations at 0.143 applying a soft-edged solvent mask. For APP/PS1 murine type III, ARTE10 murine type III, DI1, DI2 and tg-APP_Swe_ murine type II fibrils VISDEM sharpening [70] was used instead of automatic B-factor sharpening. Additional image processing information can be found in **Table S3.**

Statistics on the distribution of different polymorphs are given in **Table S2**. The term “unassigned” refers to particles that were noisy, heterogenous, and could not be used for further structure determination. Additionally, it should be noted that given statistics on polymorph distribution are based on an initial 2D classification of automatically picked particles and therefore, might be influenced by inaccurate 2D cla ssification.

#### Model Building and Refinement

For APP/PS1 murine type III fibrils and all tg-SwDI polymorphs, atomic models were built *de novo* into the computed high-resolution cryo-EM reconstructions using COOT [71]. Side chain rotamers were refined manually monitoring Ramachandran outliers and clash scores using MolProbity [72]. All models were refined using an iterative procedure of refinement in PHENIX [73] and manual modeling in COOT and ISOLDE [74].

For ARTE10 murine type III fibrils, the atomic model of APP/PS1 murine type III filaments was fitted into the density and refined using COOT, ISOLDE and PHENIX.

For ARTE10, tg-APP_Swe_, and APP23 murine type II fibrils, an atomic model of previously determined human type II Aβ filaments ([18], PDB code: 7Q4M) was fitted into the density maps and refined using COOT and PHENIX.

For tg-APP_ArcSwe_ fibrils, an atomic model of previously determined human type I Aβ filaments ([18], PDB code: 7Q4B) was fitted into the density maps and refined using COOT and PHENIX.

ChimeraX [75] was used for molecular graphics and analyses. Additional information on final models can be found in **Table S4**.

### Supplementary Tables

**Table S1:** Statistics of Data Acquisition

|  | APP/PS1 | ARTE10 | tg-SwDI | tg-APP<sub>Swe</sub> | APP23 | tg-APP<sub>ArcSwe</sub> |
| --- | --- | --- | --- | --- | --- | --- |
| Microscope | Talos Arctica | Titan Krios | Titan Krios | Titan Krios | Titan Krios | Titan Krios |
| Detector | K3 | Falcon IV | Falcon IV | Falcon IV | Falcon IV | Falcon IV |
| Energy filter slit width (e<sup>-</sup>V) | 20 | N/A | N/A | N/A | N/A | N/A |
| Magnification | 100,000 | 96,000 | 96,000 | 96,000 | 96,000 | 96,000 |
| Voltage (kV) | 200 | 300 | 300 | 300 | 300 | 300 |
| Pixel Size (Å) | 0.816 | 0.808 | 0.808 | 0.808 | 0.808 | 0.808 |
| Total electron exposure (e<sup>-</sup>/Å<sup>2</sup>) | 30.28 | 40 | 40.1 | 40.35 | 40.31 | 40.35 |
| Defocus range (μm) | [-0.5, -2.5] | [-0.5, -2.5] | [-0.5, -2.5] | [-0.5, -2.5] | [-0.5, -2.5] | [-0.5, -2.5] |
| Micrographs collected | 3674 | 9521 | 25662 | 9370 | 11622 | 17577 |

**Table S2:** Distribution of polymorphs in extracted Aβ fibril samples from different mouse models. The term “unassigned” refers to particles that were noisy, heterogenous, and could not be used for structure determination.

| Mouse Model | Distribution |
| --- | --- |
| APP/PS1 | murine type III: 100% |
| ARTE10 | murine type II: 49%; murine type III: 8%; unassigned: 43% |
| tg-SwDI | DI1: 41%; DI2: 32%; DI3: 27% |
| tg-APP<sub>Swe</sub> | murine type II: 15%; unassigned: 85% |
| APP23 | murine type II: 42%; unassigned: 58% |
| tg-APP<sub>ArcSwe</sub> | murine<sub>Arc</sub> type I: 38%; unassigned: 62% |

**Table S3:** Statistics on Image Processing

|  | **APP/PS1**<br>**murine**<br>**type III** | **ARTE10**<br>**murine**<br>**type III** | **ARTE10**<br>**murine**<br>**type II** | **tg-**<br>**SwDI**<br>**DI1** | **tg-**<br>**SwDI**<br>**DI2** | **tg-**<br>**SwDI**<br>**DI3** | **tg-APP<sup>Swe</sup>**<br>**murine**<br>**type II** | **APP23**<br>**murine**<br>**type II** | **tg-**<br>**APP<sup>ArcSwe</sup>**<br>**murine<sup>Arc</sup>**<br>**type I** |
| --- | --- | --- | --- | --- | --- | --- | --- | --- | --- |
| Final Segments (no.) | 171,432 | 10,369 | 226,920 | 54,018 | 37,937 | 10,869 | 12,100 | 119,710 | 19,036 |
| Symmetry imposed | C2 | C2 | C2 | C2 | C1 | C1 | C2 | C2 | C1 |
| Helical rise (Å) | 4.74 | 4.74 | 4.76 | 4.66 | 4.63 | 4.72 | 4.78 | 4.77 | 2.41 |
| Helical twist (°) | -1.89 | -1.91 | -3.21 | -2.78 | -2.95 | -5.24 | -3.17 | -3.23 | 179.56 |
| Map resolution (Å) | 3.5 | 3.5 | 3.4 | 3.3 | 4.2 | 4.0 | 3.8 | 3.0 | 3.0 |
| Map Sharpening B-factor | N/A | N/A | -126.1 | N/A | N/A | -90.1 | N/A | -106.5 | -80.1 |

**Table S4:**
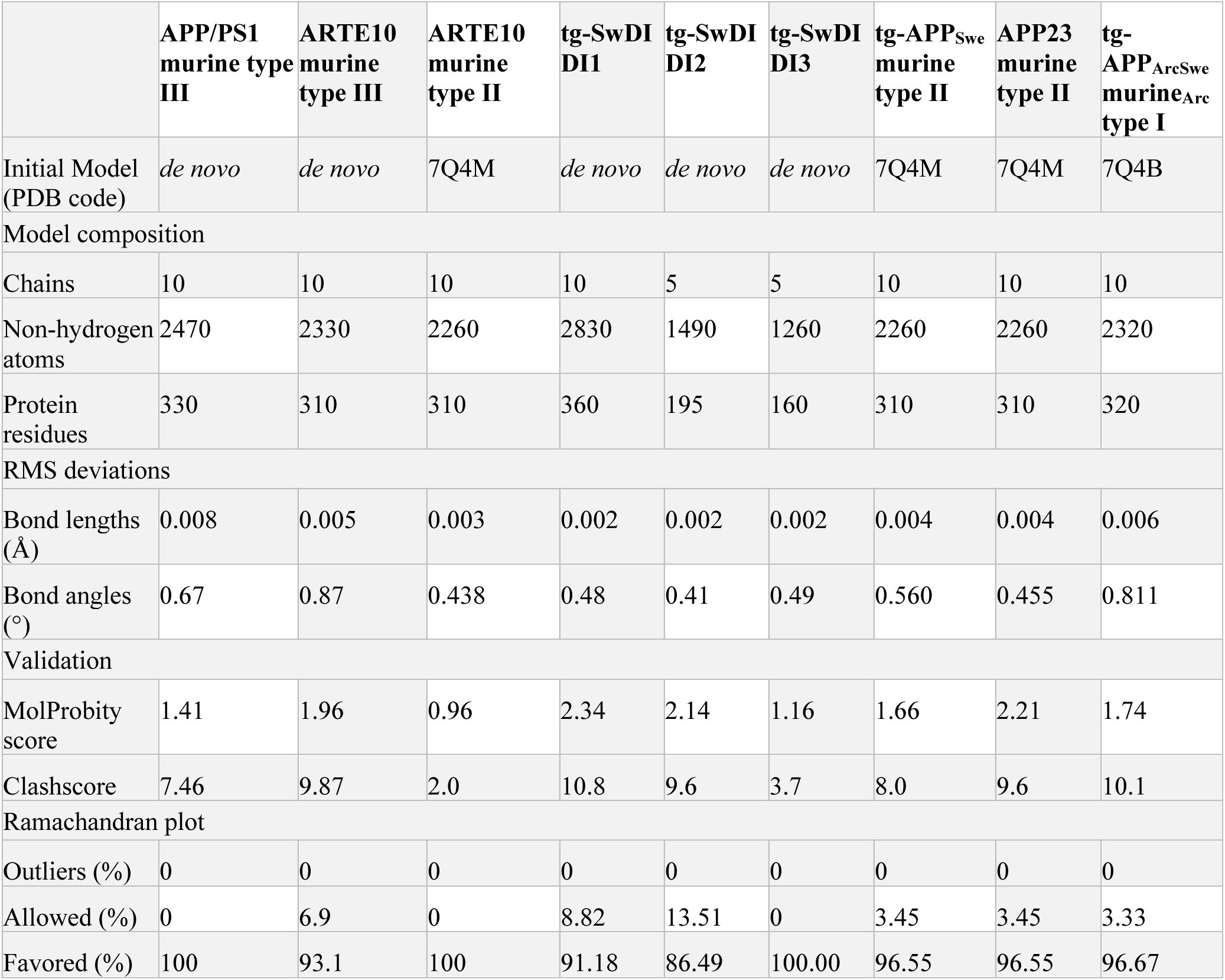
Statistics on Model Building and Refinement

### Supplementary Figures

**Figure S1:**
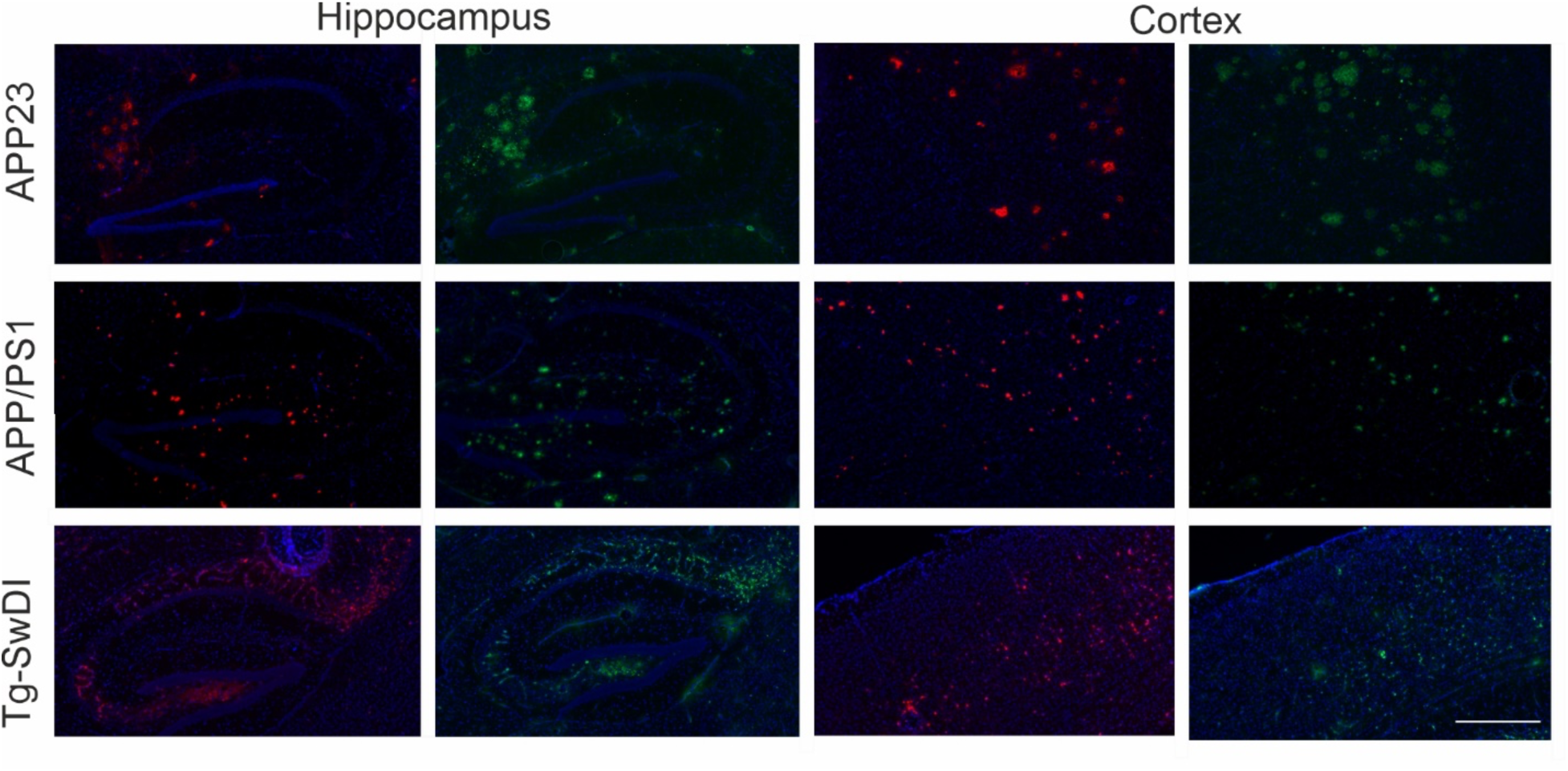
Immunohistochemical staining showing Aβ plaques in the hippocampus (left images) and cortex (right images) of APP23 (upper panel), APP/PS1 (middle panel) and tg-SwDI mice (lower panel). Two different stainings were conducted (6E10 labelled in red and 4G8 labelled in green). Nuclear staining was done with DAPI (blue). Scale bar= 500 μm.

**Figure S2:**
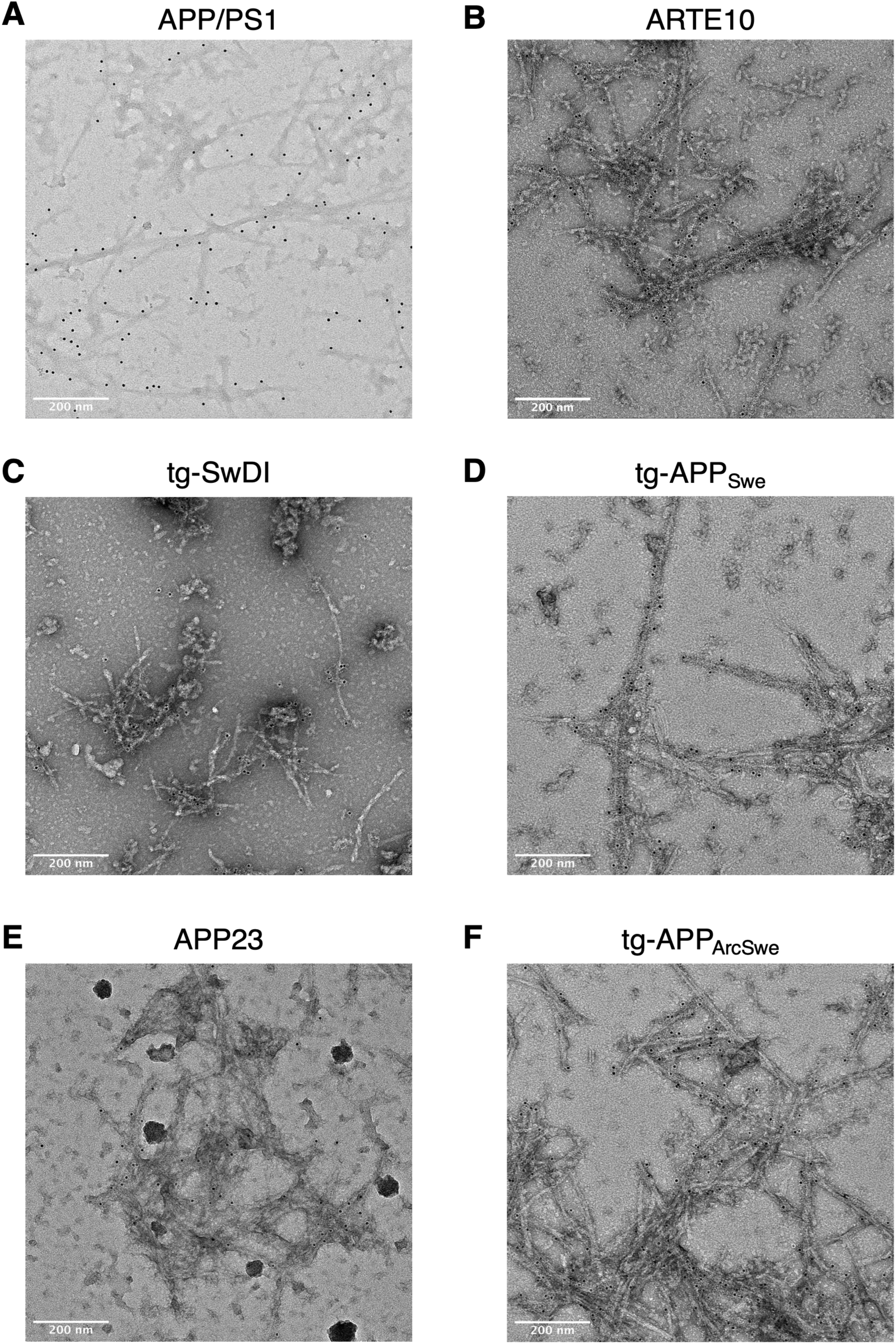
Immunogold labelling of the purified Aβ fibrils from (A) APP/PS1, (B) ARTE10, (C) tg-SwDI, (D) tg-APP_Swe_, (E) APP23 and (F) tg-APP_ArcSwe_ mouse models. NAB228 was used as primary antibody. A goat anti-mouse gold-conjugated antibody with a gold particle diameter of 6nm was used as secondary antibody.

**Figure S3:**
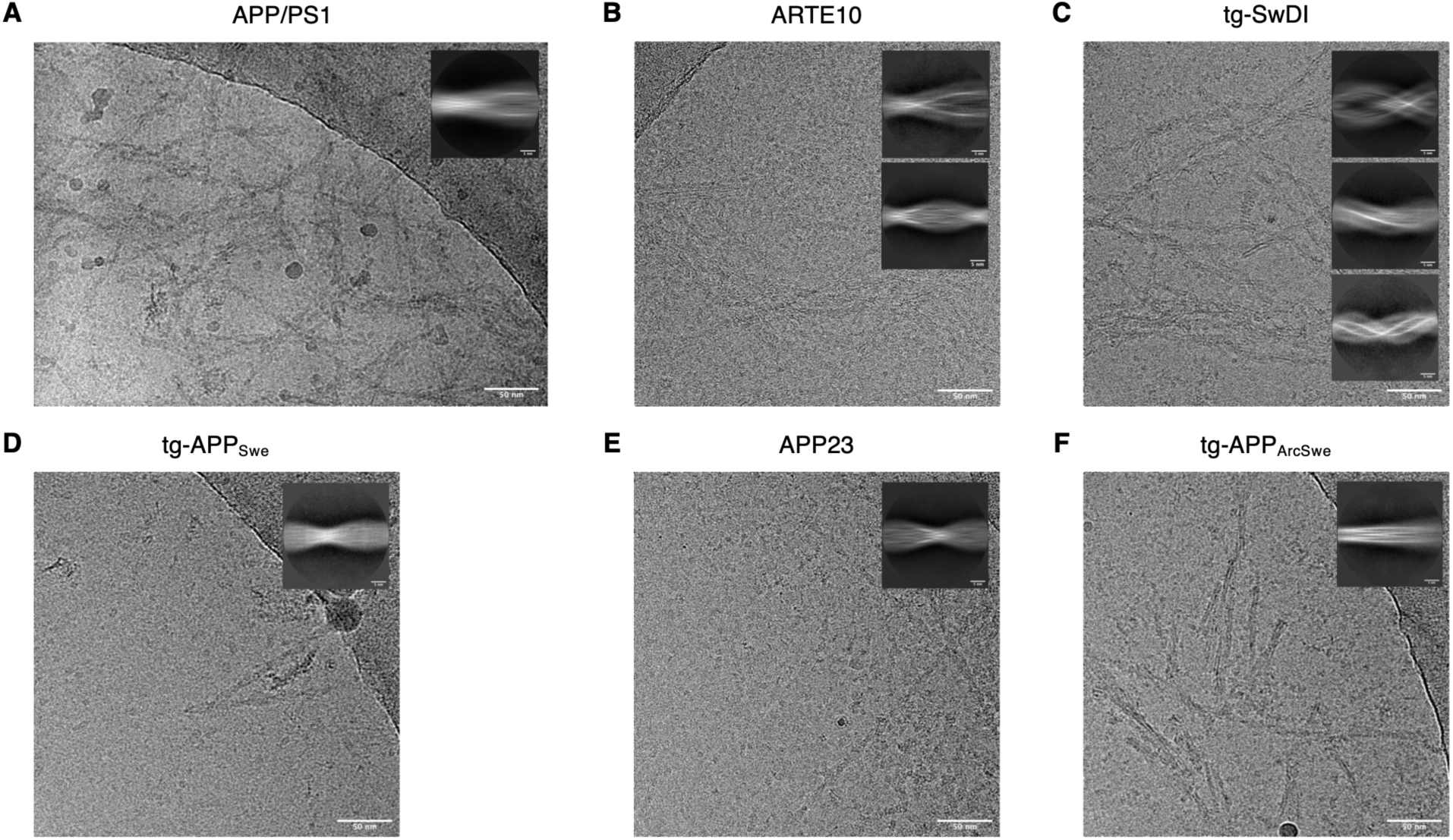
One exemplary Micrograph and 2D class of (A) APP/PS1, (B) ARTE10, (C) tg-SwDI, (D) tg-APP_Swe_, (E) APP23 and (F) tg-APP_ArcSwe_.

**Figure S4:**
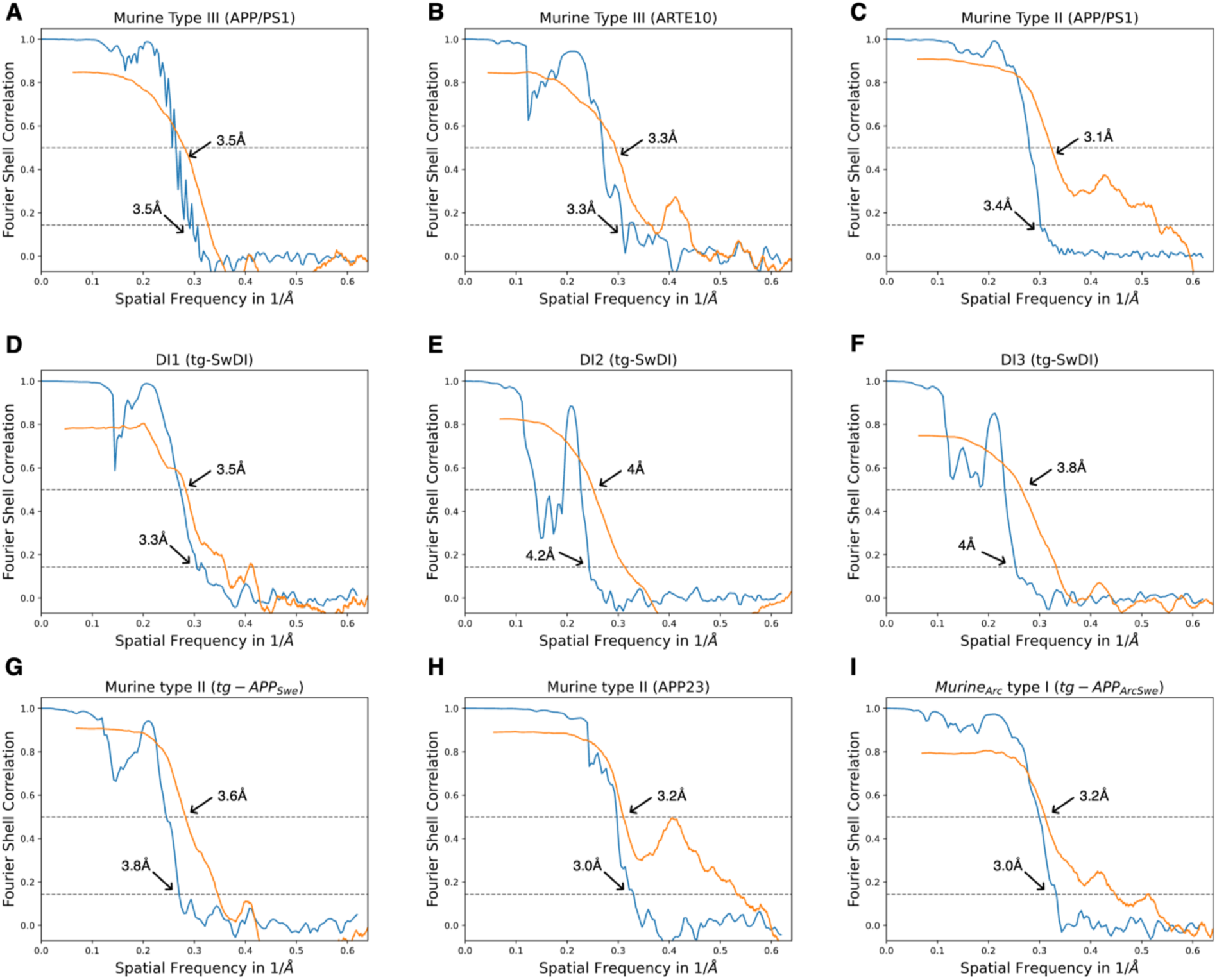
FSC curves for cryo-EM maps and structures of (A) APP/PS1 murine type III, (B) ARTE10 murine type III, (C) ARTE 10 murine type II, (D) DI1, (E) DI2, (F) DI3, (G) tg-APP_Swe_ murine type II, (H) APP23 murine type II, and (I) tg-APP_ArcSwe_ murine_Arc_ type I fibrils. FSC curves for two independently refined half maps are shown in blue; FSC curves for the refined atomic model against the final cryo-EM map in orange.

**Figure S5:**
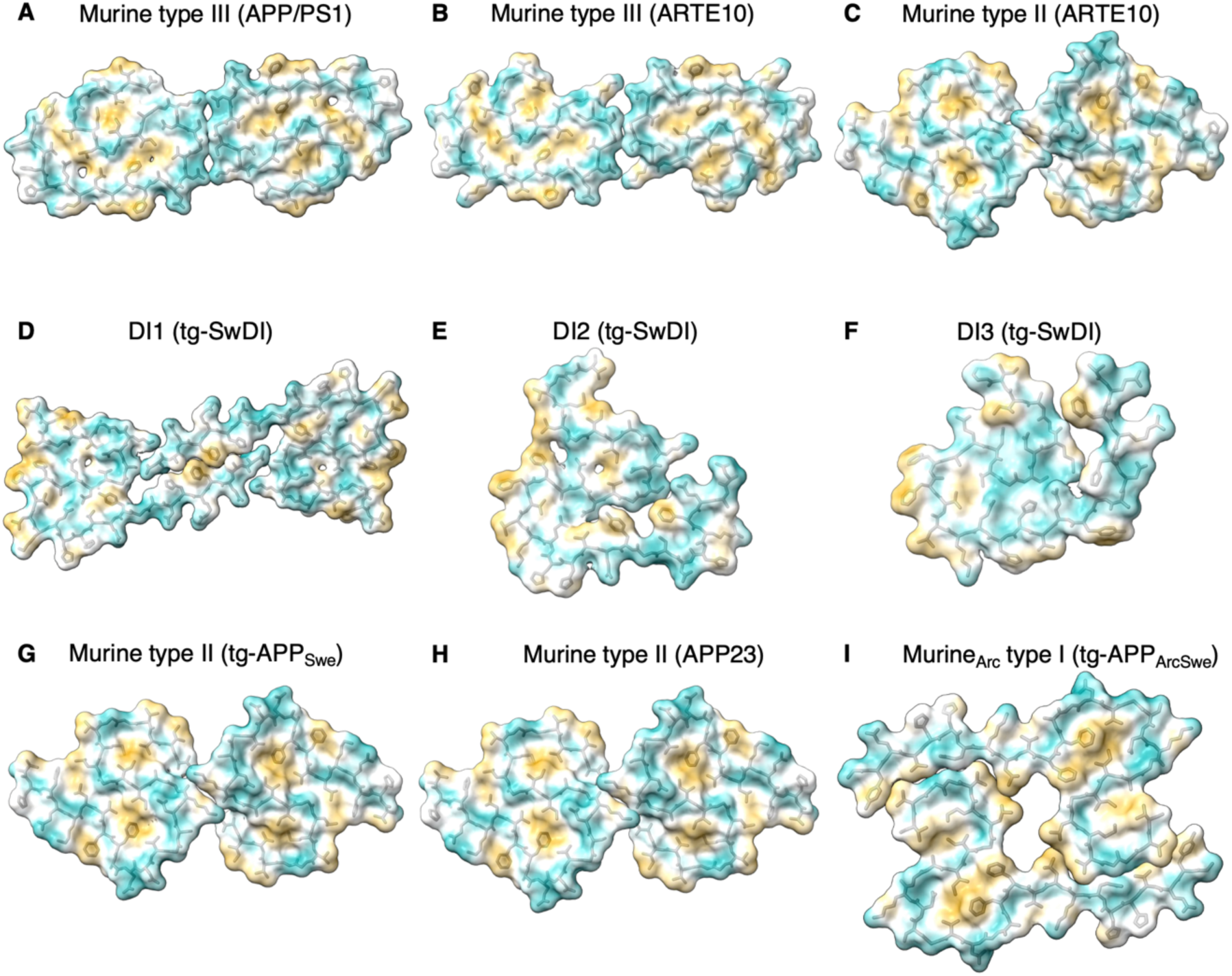
Molecular lipophilicity potential (MLP) maps for (A) APP/PS1 murine type III, (B) ARTE10 murine type III, (C) ARTE10 murine type II, (D) DI1, (E) DI2, (F) DI3, (G) tg-APP_Swe_ murine type III, (H) APP23 murine type III, and (J) murine_Arc_ type I. Coloring on the surface ranging from cyan (most hydrophilic) to white to yellow (most lipophilic/hydrophobic).

**Figure S6:**
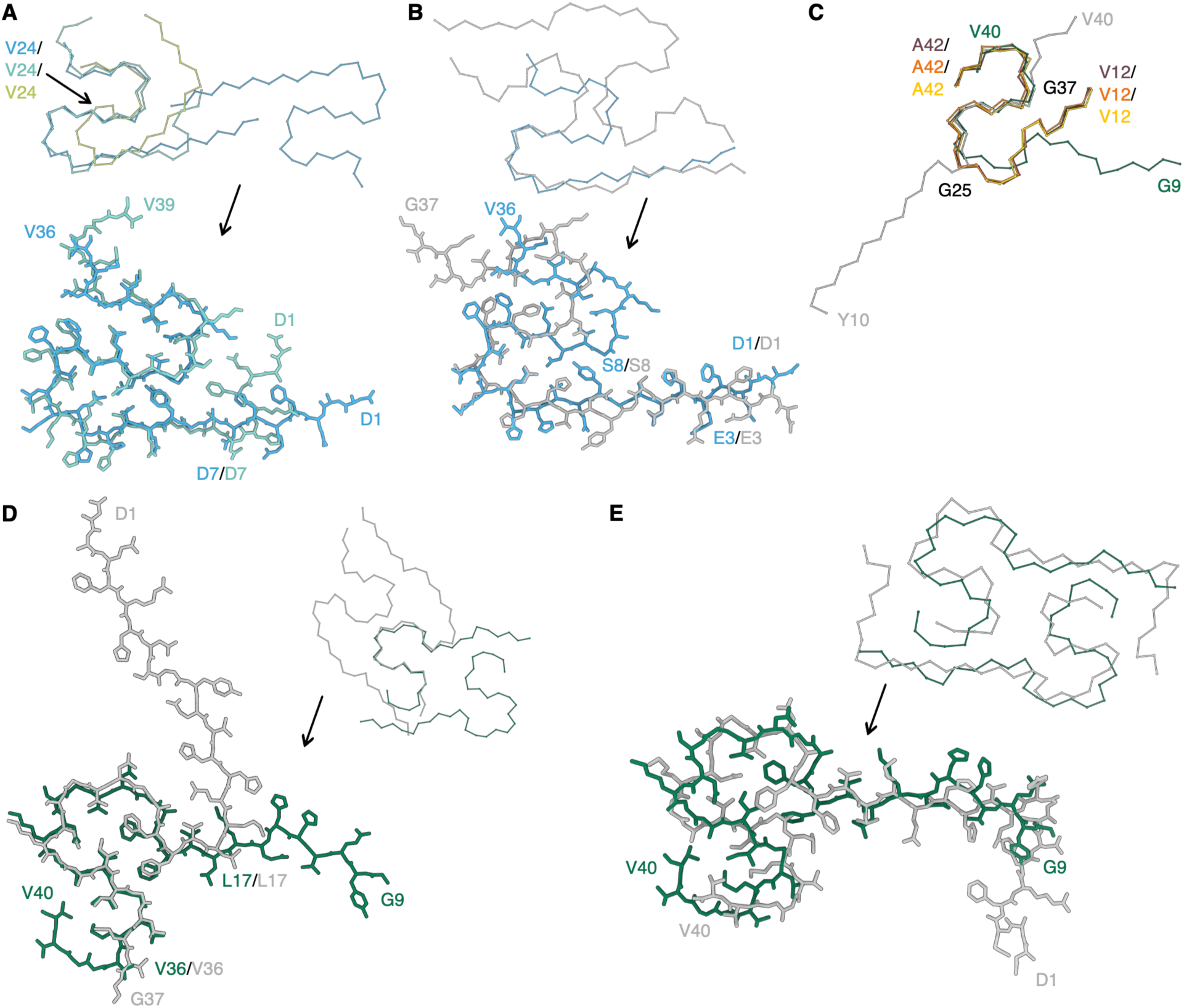
Comparison of murine Aβ fibrils with other structures. (A) Comparison of the main chain trace of DI1 (light blue) with DI2 (teal) and DI3 (light green) (top) and comparison of DI1 and DI2 (bottom). (B) Comparison of DI1 (light blue) and murine Aβ42(E22G) filaments extracted from knock-in APP^NL-G-F^ mice (gray, PDB code: 8BG9). (C) Comparison of APP23 murine type II (yellow), tg-APP_Swe_ murine type II (orange), ARTE10 murine type II (burgundy), tg-APP_ArcSwe_ murine_Arc_ type I (green) fibrils with the cryo-EM structure of Aβ40 fibrils seeded from brain homogenates from cortical tissue of an AD patient (gray, PDB code: 6W0O). (D) Comparison of tg-APP_ArcSwe_ (green) and murine Aβ42(E22G) filaments extracted from knock-in APP^NL-G-F^ mice (gray, PDB code: 8BG9). (E) Comparison of tg-APP_ArcSwe_ (green) with an NMR structure of recombinant Aβ40 E22Δ fibrils (gray, PDB code: 2MVX).

**Figure S7:**
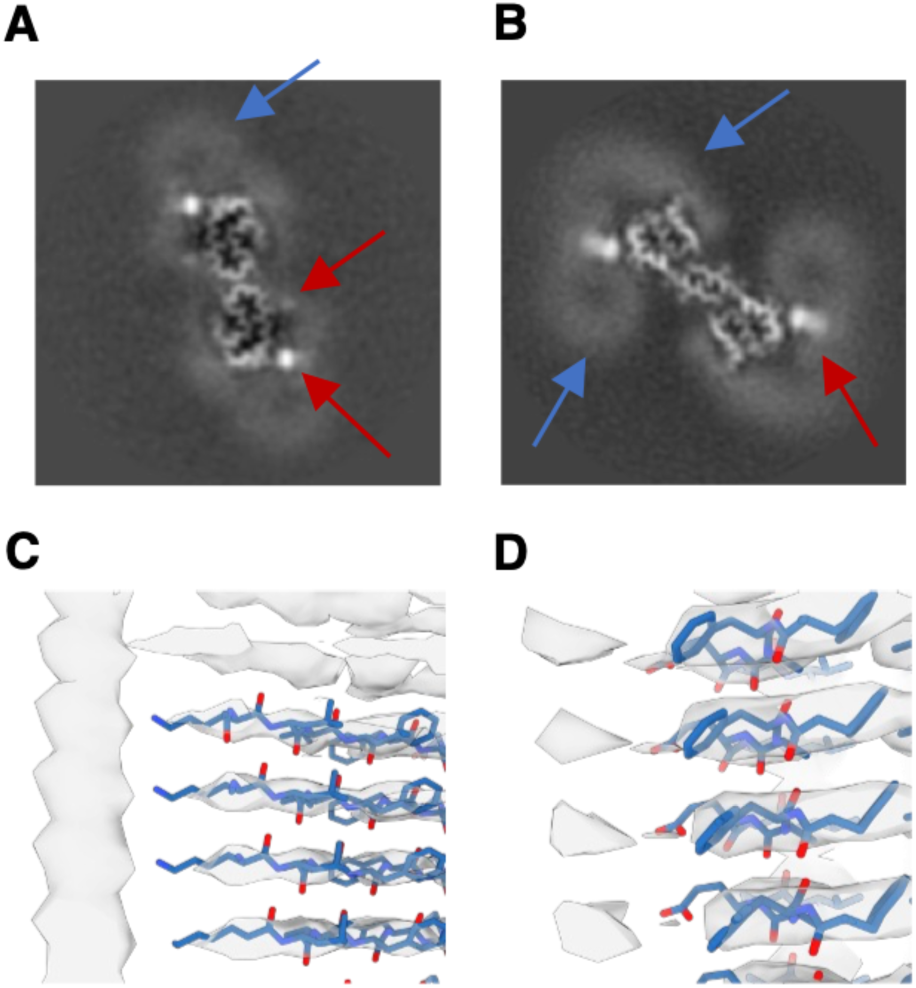
Additional densities bound to murine Aβ fibrils. (A,B) Reconstruction of Aβ fibrils extracted from (A) APP/PS1 and (B) tg-SwDI mice brain. Red arrows indicate localized, strong density, blue arrows indicate micelle-like, weak densities. (C,D) show extra densities close to (C) Lys16 and (D) Phe20/Glu22 in APP/PS1 murine type III Aβ fibrils.

## References

[1] J. M. Long and D. M. Holtzman, “Alzheimer Disease: An Update on Pathobiology and Treatment Strategies,” Cell, vol. 179, no. 2, pp. 312–339, Oct. 2019, doi: 10.1016/j.cell.2019.09.001.

[2] P. Scheltens et al., “Alzheimer’s disease,” The Lancet, vol. 388, no. 10043, pp. 505–517, Jul. 2016, doi: 10.1016/S0140-6736(15)01124-1.

[3] P. T. Nelson et al., “Correlation of Alzheimer Disease Neuropathologic Changes With Cognitive Status: A Review of the Literature,” J Neuropathol Exp Neurol, vol. 71, no. 5, pp. 362–381, May 2012, doi: 10.1097/NEN.0b013e31825018f7.

[4] F. Chiti and C. M. Dobson, “Protein Misfolding, Amyloid Formation, and Human Disease: A Summary of Progress Over the Last Decade,” Annu Rev Biochem, vol. 86, no. 1, pp. 27–68, Jun. 2017, doi: 10.1146/annurev-biochem-061516-045115.

[5] G. Chen et al., “Amyloid beta: structure, biology and structure-based therapeutic development,” Acta Pharmacol Sin, vol. 38, no. 9, pp. 1205–1235, Sep. 2017, doi: 10.1038/aps.2017.28.

[6] J. A. Hardy and G. A. Higgins, “Alzheimer’s Disease: The Amyloid Cascade Hypothesis,” Science (1979), vol. 256, no. 5054, pp. 184–185, Apr. 1992, doi: 10.1126/science.1566067.

[7] C. Zhang et al., “Amyloid-β Production Via Cleavage of Amyloid-β Protein Precursor is Modulated by Cell Density,” Journal of Alzheimer’s Disease, vol. 22, no. 2, pp. 683–694, Oct. 2010, doi: 10.3233/JAD-2010-100816.

[8] L. Gremer et al., “Fibril structure of amyloid-β(1–42) by cryo–electron microscopy,” Science (1979), vol. 358, no. 6359, pp. 116–119, 2017, doi: 10.1126/science.aao2825.

[9] L. Cerofolini et al., “Mixing Aβ(1–40) and Aβ(1–42) peptides generates unique amyloid fibrils,” Chemical Communications, vol. 56, no. 62, pp. 8830–8833, 2020, doi: 10.1039/D0CC02463E.

[10] M. T. Colvin et al., “Atomic Resolution Structure of Monomorphic Aβ42 Amyloid Fibrils,” J Am Chem Soc, vol. 138, no. 30, pp. 9663–9674, Aug. 2016, doi: 10.1021/jacs.6b05129.

[11] M. A. Wälti et al., “Atomic-resolution structure of a disease-relevant Aβ(1–42) amyloid fibril,” Proceedings of the National Academy of Sciences, vol. 113, no. 34, Aug. 2016, doi: 10.1073/pnas.1600749113.

[12] A. K. Paravastu, R. D. Leapman, W.-M. Yau, and R. Tycko, “Molecular structural basis for polymorphism in Alzheimer’s β-amyloid fibrils,” Proceedings of the National Academy of Sciences, vol. 105, no. 47, pp. 18349–18354, Nov. 2008, doi: 10.1073/pnas.0806270105.

[13] Y. Xiao et al., “Aβ(1–42) fibril structure illuminates self-recognition and replication of amyloid in Alzheimer’s disease,” Nat Struct Mol Biol, vol. 22, no. 6, pp. 499–505, Jun. 2015, doi: 10.1038/nsmb.2991.

[14] D. Willbold, B. Strodel, G. F. Schröder, W. Hoyer, and H. Heise, “Amyloid-type Protein Aggregation and Prion-like Properties of Amyloids,” Chem Rev, vol. 121, no. 13, pp. 8285–8307, Jul. 2021, doi: 10.1021/acs.chemrev.1c00196.

[15] U. Ghosh, K. R. Thurber, W.-M. Yau, and R. Tycko, “Molecular structure of a prevalent amyloid-β fibril polymorph from Alzheimer’s disease brain tissue,” Proc Natl Acad Sci U S A, vol. 118, no. 4, 2021, doi: https://doi.org/10.1073/pnas.2023089118.

[16] M. Lee, W.-M. Yau, J. M. Louis, and R. Tycko, “Structures of brain-derived 42-residue amyloid-β fibril polymorphs with unusual molecular conformations and intermolecular interactions,” Proceedings of the National Academy of Sciences, vol. 120, no. 11, Mar. 2023, doi: 10.1073/pnas.2218831120.

[17] M. Kollmer et al., “Cryo-EM structure and polymorphism of Aβ amyloid fibrils purified from Alzheimer’s brain tissue,” Nat Commun, vol. 10, no. 1, p. 4760, Dec. 2019, doi: 10.1038/s41467-019-12683-8.

[18] Y. Yang et al., “Cryo-EM structures of amyloid-β 42 filaments from human brains,” Science (1979), vol. 375, no. 6577, pp. 167–172, Jan. 2022, doi: 10.1126/science.abm7285.

[19] F. M. LaFerla and K. N. Green, “Animal Models of Alzheimer Disease,” Cold Spring Harb Perspect Med, vol. 2, no. 11, pp. a006320–a006320, Nov. 2012, doi: 10.1101/cshperspect.a006320.

[20] E. Drummond and T. Wisniewski, “Alzheimer’s disease: experimental models and reality,” Acta Neuropathol, vol. 133, no. 2, pp. 155–175, Feb. 2017, doi: 10.1007/s00401-016-1662-x.

[21] Y. Yang et al., “Cryo-EM structures of amyloid-β filaments with the Arctic mutation (E22G) from human and mouse brains,” Acta Neuropathol, vol. 145, no. 3, pp. 325–333, Mar. 2023, doi: 10.1007/s00401-022-02533-1.

[22] C. Leistner et al., “The in-tissue molecular architecture of β-amyloid in the mammalian brain,” bioRxiv, p. 2022.11.08.515609, Jan. 2022, doi: 10.1101/2022.11.08.515609.

[23] J. Cummings et al., “Alzheimer’s disease drug development pipeline: 2022,” Alzheimer’s & Dementia: Translational Research & Clinical Interventions, vol. 8, no. 1, Jan. 2022, doi: 10.1002/trc2.12295.

[24] H. Englund et al., “Sensitive ELISA detection of amyloid-β protofibrils in biological samples,” J Neurochem, vol 103, no. 1, p. 334–45, 2007

[25] D. Sehlin, X. T. Fang, L. Cato, G. Antoni, L. Lannfelt, and S. Syvänen, “Antibody-based PET imaging of amyloid beta in mouse models of Alzheimer’s disease,” Nat Commun, vol. 7, no. 1, p. 10759, Feb. 2016, doi: 10.1038/ncomms10759.

[26] A. M. Stern, et al., “Abundant Aβ fibrils in ultracentrifugal supernatants of aqueous extracts from Alzheimer’s disease brains,” bioRxiv, Oct. 2022.

[27] J. L. Cummings, T. Morstorf, and K. Zhong, “Alzheimer’s disease drug-development pipeline: few candidates, frequent failures,” Alzheimers Res Ther, vol. 6, no. 4, p. 37, 2014, doi: 10.1186/alzrt269.

[28] M. Scholl et al., “Low PiB PET retention in presence of pathologic CSF biomarkers in Arctic APP mutation carriers,” Neurology, vol. 79, no. 3, pp. 229–236, Jul. 2012, doi: 10.1212/WNL.0b013e31825fdf18.

[29] O. Philipson et al., “A highly insoluble state of Aβ similar to that of Alzheimer’s disease brain is found in Arctic APP transgenic mice,” Neurobiol Aging, vol. 30, no. 9, pp. 1393–1405, Sep. 2009, doi: 10.1016/j.neurobiolaging.2007.11.022.

[30] A. K. Schütz et al., “Atomic-Resolution Three-Dimensional Structure of Amyloid β Fibrils Bearing the Osaka Mutation,” Angewandte Chemie International Edition, vol. 54, no. 1, pp. 331–335, Jan. 2015, doi: 10.1002/anie.201408598.

[31] T. Arakhamia et al., “Posttranslational Modifications Mediate the Structural Diversity of Tauopathy Strains,” Cell, vol. 184, no. 25, pp. 6207–6210, Dec. 2021, doi: 10.1016/j.cell.2021.11.029.

[32] B. Frieg et al., “The 3D structure of lipidic fibrils of α-synuclein,” Nat Commun, vol. 13, no. 1, p. 6810, Nov. 2022, doi: 10.1038/s41467-022-34552-7.

[33] J. Davis et al., “Early-onset and Robust Cerebral Microvascular Accumulation of Amyloid β-Protein in Transgenic Mice Expressing Low Levels of a Vasculotropic Dutch/Iowa Mutant Form of Amyloid β-Protein Precursor,” Journal of Biological Chemistry, vol. 279, no. 19, pp. 20296–20306, May 2004, doi: 10.1074/jbc.M312946200.

[34] J. Miao, F. Xu, J. Davis, I. Otte-Höller, M. M. Verbeek, and W. E. Van Nostrand, “Cerebral Microvascular Amyloid β Protein Deposition Induces Vascular Degeneration and Neuroinflammation in Transgenic Mice Expressing Human Vasculotropic Mutant Amyloid β Precursor Protein,” Am J Pathol, vol. 167, no. 2, pp. 505–515, Aug. 2005, doi: 10.1016/S0002-9440(10)62993-8.

[35] G. Leinenga, W. K. Koh, and J. Götz, “A comparative study of the effects of Aducanumab and scanning ultrasound on amyloid plaques and behavior in the APP23 mouse model of Alzheimer disease,” Alzheimers Res Ther, vol. 13, no. 1, p. 76, Dec. 2021, doi: 10.1186/s13195-021-00809-4.

[36] J. Bali, A. H. Gheinani, S. Zurbriggen, and L. Rajendran, “Role of genes linked to sporadic Alzheimer’s disease risk in the production of β-amyloid peptides,” Proceedings of the National Academy of Sciences, vol. 109, no. 38, pp. 15307–15311, Sep. 2012, doi: 10.1073/pnas.1201632109.

[37] A. Lord et al., “An amyloid-β protofibril-selective antibody prevents amyloid formation in a mouse model of Alzheimer’s disease,” Neurobiol Dis, vol. 36, no. 3, pp. 425–434, Dec. 2009, doi: 10.1016/j.nbd.2009.08.007.

[38] S. Tucker et al., “The Murine Version of BAN2401 (mAb158) Selectively Reduces Amyloid-β Protofibrils in Brain and Cerebrospinal Fluid of tg-ArcSwe Mice,” Journal of Alzheimer’s Disease, vol. 43, no. 2, pp. 575–588, Nov. 2014, doi: 10.3233/JAD-140741.

[39] S. Syvänen et al., “Efficient clearance of Aβ protofibrils in AβPP-transgenic mice treated with a brain-penetrating bifunctional antibody,” Alzheimers Res Ther, vol. 10, no. 1, p. 49, Dec. 2018, doi: 10.1186/s13195-018-0377-8.

[40] L. Söderberg et al., “Lecanemab, Aducanumab, and Gantenerumab — Binding Profiles to Different Forms of Amyloid-Beta Might Explain Efficacy and Side Effects in Clinical Trials for Alzheimer’s Disease,” Neurotherapeutics, Oct. 2022, doi: 10.1007/s13311-022-01308-6.

[41] V. Logovinsky et al., “Safety and tolerability of BAN2401 - a clinical study in Alzheimer’s disease with a protofibril selective Aβ antibody,” Alzheimers Res Ther, vol. 8, no. 1, p. 14, Dec. 2016, doi: 10.1186/s13195-016-0181-2.

[42] C. J. Swanson et al., “A randomized, double-blind, phase 2b proof-of-concept clinical trial in early Alzheimer’s disease with lecanemab, an anti-Aβ protofibril antibody,” Alzheimers Res Ther, vol. 13, no. 1, p. 80, Dec. 2021, doi: 10.1186/s13195-021-00813-8.

[43] C. H. van Dyck et al., “Lecanemab in Early Alzheimer’s Disease,” New England Journal of Medicine, vol. 388, no. 1, pp. 9–21, Jan. 2023, doi: 10.1056/NEJMoa2212948.

[44] H. Kalimo et al., “The Arctic AβPP mutation leads to Alzheimer’s disease pathology with highly variable topographic deposition of differentially truncated Aβ.,” Acta Neuropathol Commun, vol. 1, p. 60, Sep. 2013, doi: 10.1186/2051-5960-1-60.

[45] O. Philipson et al., “The Arctic amyloid-β precursor protein (AβPP) mutation results in distinct plaques and accumulation of N- and C-truncated Aβ.,” Neurobiol Aging, vol. 33, no. 5, pp. 1010.e1–13, May 2012, doi: 10.1016/j.neurobiolaging.2011.10.022.

[46] A. Snellman et al., “Longitudinal Amyloid Imaging in Mouse Brain with 11C-PIB: Comparison of APP23, Tg2576, and APPswe-PS1dE9 Mouse Models of Alzheimer Disease,” Journal of Nuclear Medicine, vol. 54, no. 8, pp. 1434–1441, Aug. 2013, doi: 10.2967/jnumed.112.110163.

[47] A. Snellman et al., “In vivo PET imaging of beta-amyloid deposition in mouse models of Alzheimer’s disease with a high specific activity PET imaging agent [18F]flutemetamol,” EJNMMI Res, vol. 4, no. 1, p. 37, Dec. 2014, doi: 10.1186/s13550-014-0037-3.

[48] B. H. Yousefi et al., “FIBT versus florbetaben and PiB: a preclinical comparison study with amyloid-PET in transgenic mice,” EJNMMI Res, vol. 5, no. 1, p. 20, Dec. 2015, doi: 10.1186/s13550-015-0090-6.

[49] A. Willuweit et al., “Comparison of the Amyloid Load in the Brains of Two Transgenic Alzheimer’s Disease Mouse Models Quantified by Florbetaben Positron Emission Tomography,” Front Neurosci, vol. 15, Oct. 2021, doi: 10.3389/fnins.2021.699926.

[50] S. R. Meier et al., “11C-PiB and 124 I-Antibody PET Provide Differing Estimates of Brain Amyloid-β After Therapeutic Intervention,” Journal of Nuclear Medicine, vol. 63, no. 2, pp. 302–309, Feb. 2022, doi: 10.2967/jnumed.121.262083.

[51] Takanori Nakane, “atom2svg.” 2020. Accessed: May 01, 2022. [Online]. Available: https://zenodo.org/record/4090925#.YqNEES-21iM

[52] J. L. Jankowsky et al., “Mutant presenilins specifically elevate the levels of the 42 residue β-amyloid peptide in vivo: evidence for augmentation of a 42-specific γ secretase,” Hum Mol Genet, vol. 13, no. 2, pp. 159–170, Jan. 2004, doi: 10.1093/hmg/ddh019.

[53] C. Janus, A. Y. Flores, G. Xu, and D. R. Borchelt, “Behavioral abnormalities in APPSwe/PS1dE9 mouse model of AD-like pathology: comparative analysis across multiple behavioral domains,” Neurobiol Aging, vol. 36, no. 9, pp. 2519–2532, Sep. 2015, doi: 10.1016/j.neurobiolaging.2015.05.010.

[54] M. Garcia-Alloza et al., “Characterization of amyloid deposition in the APPswe/PS1dE9 mouse model of Alzheimer disease,” Neurobiol Dis, vol. 24, no. 3, pp. 516–524, Dec. 2006, doi: 10.1016/j.nbd.2006.08.017.

[55] A. Willuweit et al., “Early-Onset and Robust Amyloid Pathology in a New Homozygous Mouse Model of Alzheimer’s Disease,” PLoS One, vol. 4, no. 11, p. e7931, Nov. 2009, doi: 10.1371/journal.pone.0007931.

[56] F. Xu et al., “Early-onset subicular microvascular amyloid and neuroinflammation correlate with behavioral deficits in vasculotropic mutant amyloid β-protein precursor transgenic mice,” Neuroscience, vol. 146, no. 1, pp. 98–107, Apr. 2007, doi: 10.1016/j.neuroscience.2007.01.043.

[57] J. Davis et al., “Early-onset and Robust Cerebral Microvascular Accumulation of Amyloid β-Protein in Transgenic Mice Expressing Low Levels of a Vasculotropic Dutch/Iowa Mutant Form of Amyloid β-Protein Precursor,” Journal of Biological Chemistry, vol. 279, no. 19, pp. 20296–20306, May 2004, doi: 10.1074/jbc.M312946200.

[58] A. Lord, H. Kalimo, C. Eckman, X.-Q. Zhang, L. Lannfelt, and L. N. G. Nilsson, “The Arctic Alzheimer mutation facilitates early intraneuronal Aβ aggregation and senile plaque formation in transgenic mice,” Neurobiol Aging, vol. 27, no. 1, pp. 67–77, Jan. 2006, doi: 10.1016/j.neurobiolaging.2004.12.007.

[59] S. Lillehaug, G. H. Syverstad, L. N. G. Nilsson, J. G. Bjaalie, T. B. Leergaard, and R. Torp, “Brainwide distribution and variance of amyloid-beta deposits in tg-ArcSwe mice,” Neurobiol Aging, vol. 35, no. 3, pp. 556–564, Mar. 2014, doi: 10.1016/j.neurobiolaging.2013.09.013.

[60] W. Michno et al., “Pyroglutamation of amyloid-βx-42 (Aβx-42) followed by Aβ1–40 deposition underlies plaque polymorphism in progressing Alzheimer’s disease pathology,” Journal of Biological Chemistry, vol. 294, no. 17, pp. 6719–6732, Apr. 2019, doi: 10.1074/jbc.RA118.006604.

[61] N. M. Gulati, U. Torian, J. R. Gallagher, and A. K. Harris, “Immunoelectron Microscopy of Viral Antigens,” Curr Protoc Microbiol, vol. 53, no. 1, Jun. 2019, doi: 10.1002/cpmc.86.

[62] D. Tegunov and P. Cramer, “Real-time cryo-electron microscopy data preprocessing with Warp,” Nat Methods, vol. 16, no. 11, pp. 1146–1152, Nov. 2019, doi: 10.1038/s41592-019-0580-y.

[63] A. Rohou and N. Grigorieff, “CTFFIND4: Fast and accurate defocus estimation from electron micrographs,” J Struct Biol, vol. 192, no. 2, pp. 216–221, Nov. 2015, doi: 10.1016/j.jsb.2015.08.008.

[64] S. He and S. H. W. Scheres, “Helical reconstruction in RELION,” J Struct Biol, vol. 198, no. 3, pp. 163–176, 2017, doi: 10.1016/j.jsb.2017.02.003.

[65] J. Zivanov et al., “New tools for automated high-resolution cryo-EM structure determination in RELION-3,” Elife, vol. 7, Nov. 2018, doi: 10.7554/eLife.42166.

[66] T. Wagner, L. Lusnig, S. Pospich, M. Stabrin, F. Schönfeld, and S. Raunser, “Two particle-picking procedures for filamentous proteins: SPHIRE-crYOLO filament mode and SPHIRE-STRIPER,” Acta Crystallogr D Struct Biol, vol. 76, no. 7, pp. 613–620, Jul. 2020, doi: 10.1107/S2059798320007342.

[67] T. Wagner et al., “SPHIRE-crYOLO is a fast and accurate fully automated particle picker for cryo-EM,” Commun Biol, vol. 2, no. 1, p. 218, Jun. 2019, doi: 10.1038/s42003-019-0437-z.

[68] X. Bai, E. Rajendra, G. Yang, Y. Shi, and S. H. Scheres, “Sampling the conformational space of the catalytic subunit of human γ-secretase,” Elife, vol. 4, Dec. 2015, doi: 10.7554/eLife.11182.

[69] S. H. W. Scheres, “Amyloid structure determination in RELION-3.1,” Acta Crystallogr D Struct Biol, 2020, doi: 10.1107/S2059798319016577.

[70] M. Spiegel, A. K. Duraisamy, and G. F. Schröder, “Improving the visualization of cryo-EM density reconstructions,” J Struct Biol, vol. 191, no. 2, pp. 207–213, Aug. 2015, doi: 10.1016/j.jsb.2015.06.007.

[71] A. Casañal, B. Lohkamp, and P. Emsley, “Current developments in Coot for macromolecular model building of Electron Cryo-microscopy and Crystallographic Data,” Protein Science, vol. 29, no. 4, pp. 1055–1064, Apr. 2020, doi: 10.1002/pro.3791.

[72] C. J. Williams et al., “MolProbity: More and better reference data for improved all-atom structure validation,” Protein Science, vol. 27, no. 1, pp. 293–315, Jan. 2018, doi: 10.1002/pro.3330.

[73] P. V. Afonine et al., “Real-space refinement in PHENIX for cryo-EM and crystallography,” Acta Crystallogr D Struct Biol, vol. 74, no. 6, pp. 531–544, Jun. 2018, doi: 10.1107/S2059798318006551.

[74] T. I. Croll, “ISOLDE: a physically realistic environment for model building into low-resolution electron-density maps,” Acta Crystallogr D Struct Biol, vol. 74, no. 6, pp. 519–530, Jun. 2018, doi: 10.1107/S2059798318002425.

[75] E. F. Pettersen et al., “UCSF ChimeraX : Structure visualization for researchers, educators, and developers,” Protein Science, vol. 30, no. 1, pp. 70–82, Jan. 2021, doi: 10.1002/pro.3943.

